# CynoBrain: A unified high-resolution framework for cross-modal data integration for macaque brain mapping

**DOI:** 10.64898/2026.09.07.749786

**Authors:** Hao Yang, MingLi Wang, NiNi Yuan, Mengchao Pei, Kaiwei Zhang, Binshi Bo, Yin Xia, ZhiMing Shen, Zhifeng Liang

## Abstract

The cynomolgus macaque is a widely used primate model in neuroscience, yet existing brain atlas resources remain limited in resolution and lack a standardized framework for integrating structural, molecular, and connectivity data within a common reference space. Here we present CynoBrain, a high-resolution multi-modal brain atlas constructed from population-averaged MRI data at 9.4 Tesla, yielding a 75 µm isotropic template with significant improvement in resolution. A co-registered whole-brain autofluorescence template resolves cytoarchitectural boundaries not visible in MRI template. Three major parcellation schemes, D99 143-area, M132 105-area, and SARM are incorporated in CynoBrain space. Spatial transcriptomic based laminar definition, reconstructed from 2D sections into volumetric common space, provides a six-layer cortical segmentation spanning the whole cortex. As a demonstration of the integrative capability of this framework, 2D retrograde tracing data is reconstructed and mapped to the Cynobrain space, enabling volumetric compilation of 2D macaque brain mapping data across subjects. An online interactive platform was constructed to allow full access to Cynobrain, establishing it as a scalable, open platform for cross-modal data integration for primate neuroscience. Overall, CynoBrain is a unified framework for cross-modal data integration for macaque brain mapping with high-resolution, multiple atlases and transcriptomic based laminar definition.

## 1. Introduction

Non-human primates play an important role in modern biomedical research, with macaques emerging as the preferred experimental model. Macaques share approximately 93% of their genomic sequence with humans[1, 2], making them ideal models for investigating human disease mechanisms and developing therapeutic approaches. The neural system of macaques closely resembles that of humans, including a well-developed prefrontal cortex, complex cortical laminar organization, and comparable neural circuit organization and connectivity patterns [3–5].

These similarities have motivated a broad range of data-intensive research efforts — each generating rich but independently organized datasets over the past several decades. Classically, neuroanatomical tracing studies have generated extensive connectivity data in the macaque brain, mapping detailed circuit diagrams through both traditional and modern viral tracing techniques. In particular, the landmark retrograde tracing program by Markov has systematically quantified inter-areal connection weights across 91 cortical areas, producing the most comprehensive directed connectivity matrix available for any primate brain [6]. Technical advances have recently produced single-neuron level projectome in macaque brain [7]. Macaque spatial transcriptomic data has also seen remarkable growth through large-scale initiatives such as the NIH Blueprint Non-Human Primate Atlas[8] and the China Brain Project [9–11] .

This breadth of the above multimodal data, however, presents a significant challenge for data integration in space. Currently, most studies either present the data in native space [12–14] or reduce the space information to brain region level. The latter approach leads to the dependency on parcellation schemes, which are highly variable and without a consensus. Therefore, much of the newer data remains scattered across individual studies and are difficult to integrate. The CoCoMac database[15, 16] represented an early attempt to collate connectivity data from various tracing studies, yet its coverage remains limited to a subset of published connections[17] and does not incorporate the quantitative weight information provided by more recent studies[18, 19].

To be able to fully integrate multimodal brain mapping data in macaque, brain atlases, as well as pipelines that map data to atlas space, are essential. Traditional macaque brain atlases include Paxinos atlas[20] and the D99 atlas[21], with newer INIA19 template [22], Yerkes19 [23–25], and the NMT [26] representing significant advances by providing population-averaged MRI reference spaces. CHARM and SARM atlases [27] further expanded these resources by offering six-level hierarchical probabilistic maps of cortical and subcortical regions.

Despite these progress, current atlases still face critical limitations that prevent comprehensive cross-modal integration. First, the resolution of existing macaque brain atlases remains insufficient to resolve fine-grained cortical and subcortical structures. Newer data, such as spatial transcriptomic or single-neuron level projectome data are natively ultrahigh resolution, and such mismatch between resolution in raw data and brain atlases creates barriers for data integration. Particularly, cortical layer definition has not been integrated into macaque brain atlases yet, significantly lagging mouse atlas [28]. Cortical layers are a fundamental feature of mammalian cortex, thus providing layer definition in a standard reference space is highly beneficial when integrating multimodal data (e.g., single-neuron projection data). Lastly, as a large number of data modalities are in 2D space, a robust computational pipeline is required to reconstruct those 2D data into 3D atlas space, therefore enabling a seamless data integration across subjects and modalities.

To this end, we developed CynoBrain, a high-resolution cynomolgus macaque brain template and comprehensive multimodal integration platform. This platform provides a standardized workflow for registering and integrating diverse data types into a unified reference framework, including anatomical, molecular, and connectivity information. Built upon a population-averaged 9.4 T MRI template at 75 µm isotropic resolution, CynoBrain incorporates a co-registered whole-brain autofluorescence template. Three parcellation schemes (D99 143-area, M132 105-area, and SARM) and transcriptome-derived six-layer cortical laminar maps are all incorporated in the same CynoBrain reference space. To demonstrate the utility of this platform, 2D retrograde tracing data in two macaques were reconstructed and mapped into the CynoBrain reference space, enabling direct 3D data atlasing of conventional 2D data. Finally, an interactive online platform was constructed to allow full access to CynoBrain, a scalable, open resource platform for cross-modal data integration for primate neuroscience.

## 2. Results

### 2.1. A High-Resolution Multimodal Template of the Cynomolgus Macaque Brain

We constructed the CynoBrain template from population-averaged ex vivo MRI data acquired from 22 cynomolgus macaques (Supplementary Table 1) at 9.4 Tesla, yielding a T2*-weighted volumetric reference at 75 µm isotropic resolution. The template provides consistent tissue contrast and geometric fidelity across all standard viewing planes, capturing both gross morphological features and fine structural details across the entire brain (Fig. 1A).

**Figure 1.**
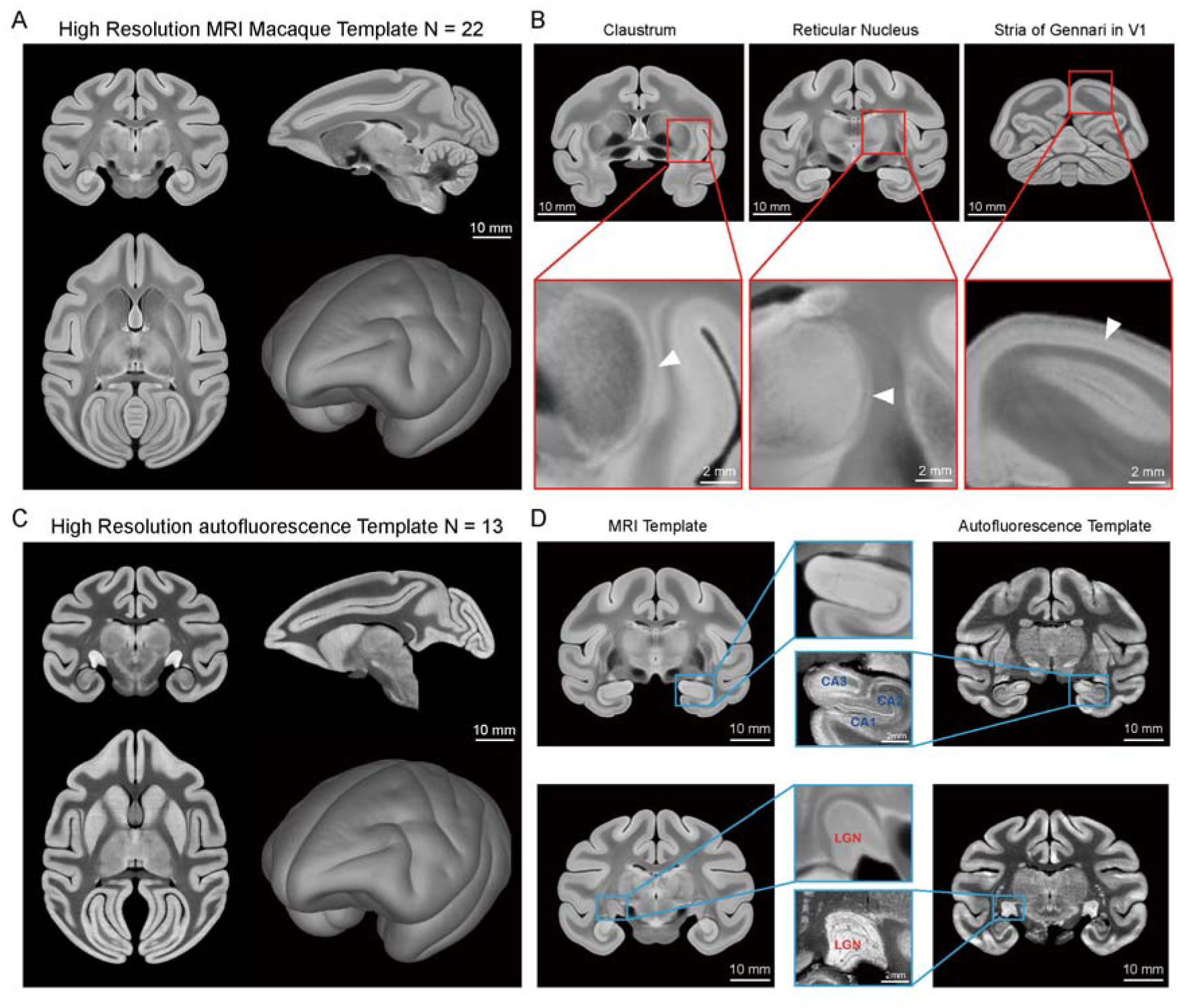
High-resolution population-averaged MRI and autofluorescence templates of the macaque brain. **A** Population-averaged T2*-weighted MRI template from 22 macaque brains shown in coronal, sagittal, axial, and lateral views. **B** Magnified views demonstrating fine anatomical structures including claustrum (left, arrowhead), reticular nucleus of thalamus (middle, arrowhead), and Stria of Gennari in V1 (right, arrowhead). **C** Population-averaged autofluorescence template from 12 macaque brains acquired via fluorescence sections. **D** Multimodal comparison showing hippocampal subfields (CA1, CA2, CA3) and lateral geniculate nucleus (LGN) with different contrast in fluorescence versus MRI templates.

The resolving power of the CynoBrain template was assessed by examining three anatomical structures that serve as demanding benchmarks for MRI template quality (Fig. 1B). The reticular nucleus of the thalamus, a thin laminar gray matter sheet lining the lateral and anterior aspects of the thalamus, was resolved as a continuous band distinguishable from the internal capsule and adjacent thalamic nuclei. The claustrum, situated between the insular cortex and the putamen, was clearly delineated as a distinct thin sheet of gray matter separated from adjacent white matter. Lastly, the Stria of Gennari within primary visual cortex (V1), corresponding to the dense band of myelinated fibers in layer IV, was visible as a hyperintense stripe within the cortical ribbon.

Furthermore, a population-averaged autofluorescence template was constructed from 12 macaque brains using whole-brain autofluorescence sections (see Methods) (Fig. 1C) to complement the MRI template. Serial two-dimensional autofluorescence images underwent blockface-guided registration to establish an initial volumetric reconstruction, which was subsequently co-registered to the reference template via the corresponding ex vivo MRI volume (Supplementary Fig. 1 and 2). Multimodal comparison demonstrated that the two templates provide complementary anatomical information in regions where one modality alone is insufficient for reliable boundary delineation. Within the hippocampal formation, the subfields CA1, CA2, and CA3 were clearly distinguishable in the autofluorescence template based on differences in laminar organization, whereas the corresponding MRI template showed insufficient contrast. Similarly, the laminar structure of the lateral geniculate nucleus (LGN) was sharply delineated in the autofluorescence template (Fig. 1D). These results demonstrate that the two modalities provide complementary contrast information, with MRI offering volumetric coverage and geometric standardization, and autofluorescence imaging provided different tissue contrast.

**Figure 2.**
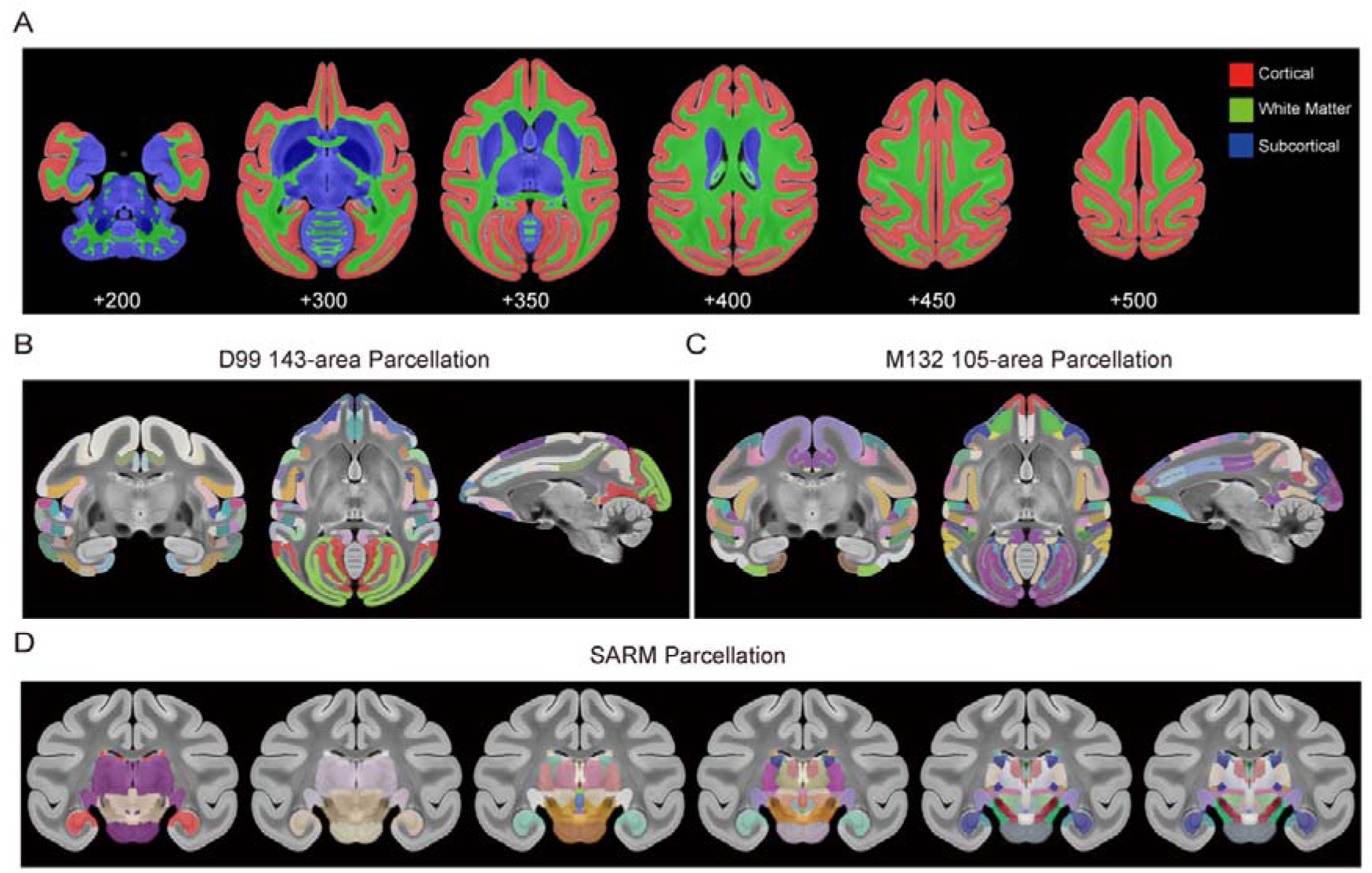
Anatomical parcellation schemes for the macaque brain template. **A** Three-class tissue segmentation showing cortical gray matter (red), subcortical gray matter (blue), and white matter (green) across six representative axial slices from ventral to dorsal levels. **B-C** Cortical parcellation schemes: D99 atlas with 143 cortical and subcortical areas (left) and M132 105-area cortical parcellation (right), displayed in coronal, axial, and lateral views overlaid on the template. **D** SARM subcortical parcellation showing detailed segmentation of subcortical structures including thalamic nuclei, basal ganglia, hippocampus, and amygdala across six coronal slices from anterior to posterior levels.

### 2.2. Comprehensive Parcellation of Cortical and Subcortical Structures

To support quantitative region-of-interest analyses and cross-modal data integration within the CynoBrain framework, we implemented a multi-atlas parcellation approach covering cortical, subcortical, and white matter compartments. A three-class tissue segmentation of the CynoBrain template produced tissue masks for cortical gray matter, subcortical gray matter, and white matter across the full anterior-to-posterior extent of the brain (Fig. 2A).

To enable region-specific analysis across the full extent of the CynoBrain atlas, we incorporated both cortical and subcortical parcellation schemes into a common volumetric space. For cortical parcellation, two complementary schemes were registered into CynoBrain space. A subset of D99 atlas[27] defined 143 cortical regions[12] in NMT v2 space (Fig. 2B). The M132 parcellation, comprising 105 cortical areas and originally defined exclusively on a two-dimensional surface mesh, was projected into three-dimensional volumetric space using a surface-informed label optimization procedure (see Methods). Label boundaries were verified to remain perpendicular to the cortical surface throughout the full depth of the cortical ribbon, ensuring geometric fidelity of the reconstructed parcellation (Fig. 2C). This volumetric reconstruction of M132 parcellation enables its direct application in neuroimaging datasets.

Subcortical structures were delineated using the SARM atlas, which provided parcellations of individual thalamic nuclei, basal ganglia components, hippocampal subfields, and amygdaloid nuclei (Fig. 2D). Together, the cortical and subcortical parcellations establish a comprehensive, multi-level reference framework within CynoBrain space, supporting both region-of-interest analyses and whole-brain connectivity studies.

### 2.3. Transcriptomic-defined Laminar Architecture of macaque cortex delineated in CyoBrain space

Building upon our previously established whole-brain transcriptome macaque atlas[12], we sought to integrate the transcriptomic-defined laminar organization of the cerebral cortex into the CynoBrain MRI template. The cortical layers were defined by transcriptomic profiling of total RNA sections combined with adjacent NeuN staining sections (Supplementary Fig. 3). A computational registration pipeline (see Methods) was then employed to project these two-dimensional laminar definitions into volumetric reference space (Fig. 3). This pipeline incorporated slice-to-volume registration and vertex Laplacian smoothing for cortical surface reconstruction, generating a layer-specific cortical segments anchored to the template.

**Figure 3.**
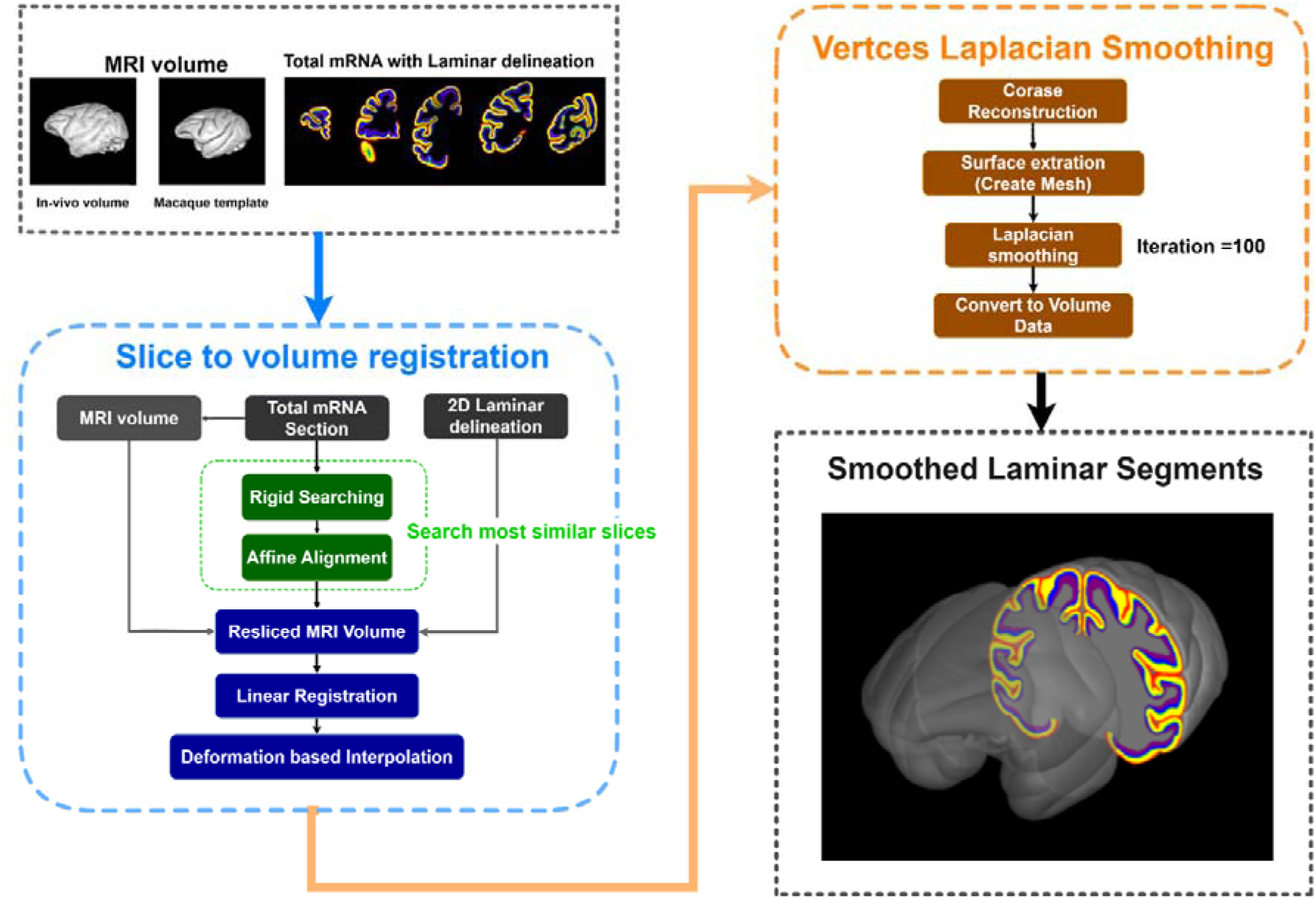
Pipeline for transcriptome-derived cortical layers reconstruction. Workflow for mapping transcriptome-defined cortical layers to the MRI template: slice-to-volume registration of total RNA sections (blue box) and vertex Laplacian smoothing for cortical surface reconstruction (orange box), generating layer-specific cortical parcellation on the template.

The resulting layer-specific parcellation was visualized across the anterior-to-posterior extent of the cortex in a 3D rendering (Fig. 4A, top pannel). Two representative coronal sections were selected to highlight the detailed laminar architecture within specific regions (Fig. 4A, bottom pannel): left spanel illustrating the parietal cortex with clearly delineated LIPd, LIPv, and area 7a/7b boundaries, and right posterior slices demonstrating the insula and somatosensory cortex, including areas Id and SII.

**Figure 4.**
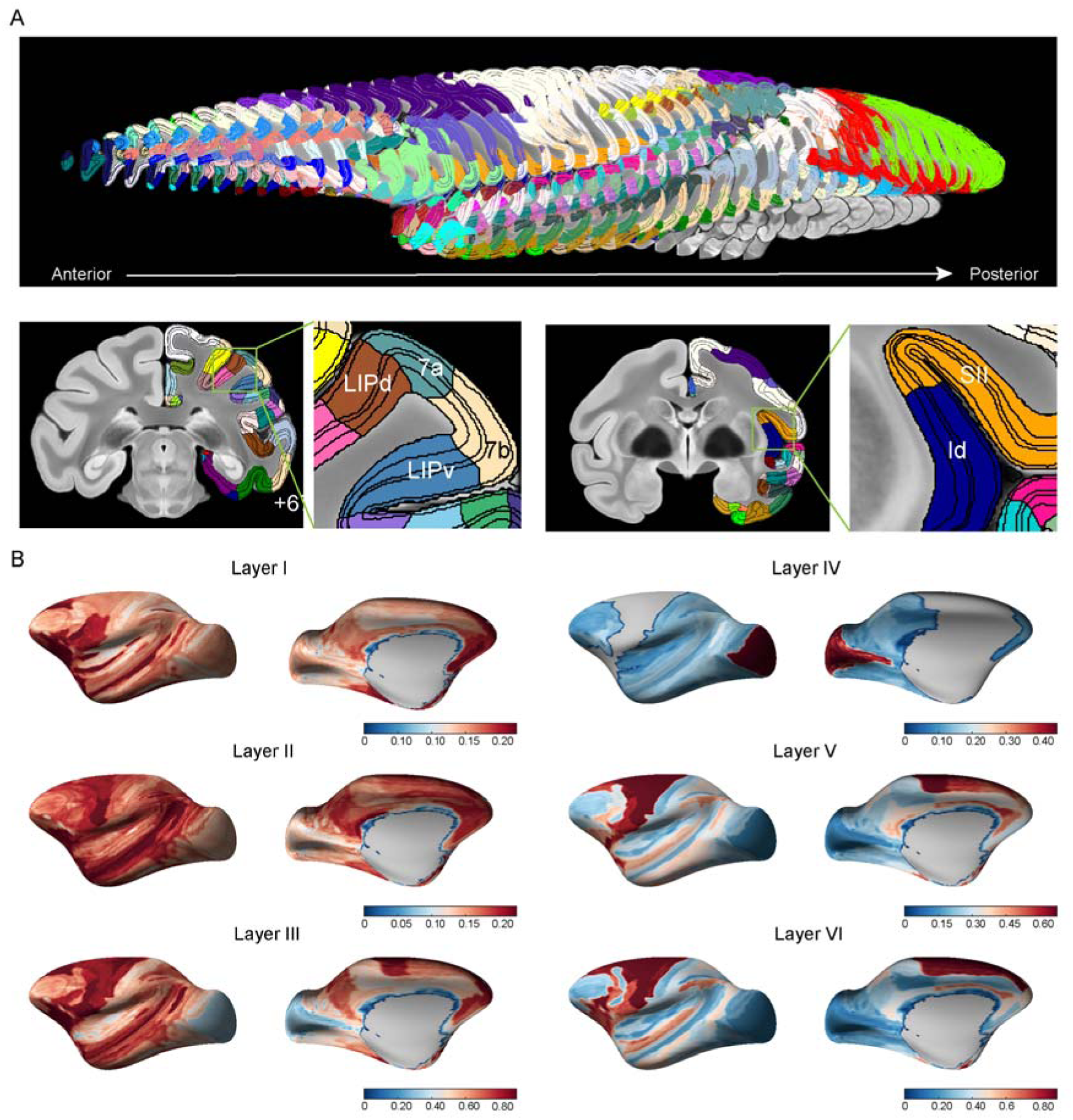
Transcriptome-derived laminar organization and thickness profiles across cortex. **A** Transcriptome-derived layer segmentation illustrating laminar organization across cortical regions (top). Two representative slices are zoomed in to highlight the detailed laminar architecture (bottom). **B** Cortical thickness profiles across brain regions (prefrontal, frontal, cingulate, somatosensory, parietal, temporal, auditory and occipital cortices) visualized on the cortical surface.

To quantify regional patterns of cortical laminar architecture, we measured layer-specific thickness from the native 2D sections across 143 cortical regions (Fig. 4B and Supplementary Table 2). Layer III exhibited the greatest mean thickness (0.692 ± 0.165 mm) and was the dominant layer across most cortical areas. Layer VI showed substantial thickness (0.581 ± 0.263 mm), particularly prominent in limbic and prefrontal regions. Layer V was moderately thick (0.415 ± 0.139 mm), especially in motor cortex. Layer I (0.210 ± 0.056 mm) and Layer II (0.194 ± 0.029 mm) were relatively thin and uniform across regions. Layer IV was thinnest (0.105 ± 0.083 mm) and absent in agranular cortex regions. The primary visual cortex (V1) displayed a distinct pattern with Layer IV as the thickest layer (0.506 mm), consistent with its granular architecture. Motor cortex (F1, F2, F7) showed prominent Layer V (0.757 mm) and Layer VI (1.115 mm), reflecting its role in motor output. Prefrontal cortex exhibited thicker Layer III (0.772 mm) compared to visual cortex (0.626 mm), suggesting enhanced cortico-cortical connectivity in association areas.

### 2.4 Mapping Retrograde Tracing Data to the CynoBrain Reference Space

Having established the framework of CynoBrain reference space, we next used this framework to reconstruct retrogradely labeled neurons from 2D slices to our 3D reference space. Retrograde tracers were injected to two cortical areas: area 23 and claustrum (Supplementary Fig. 4 and 5). The 2D autofluorescence images overlaid with labeled neurons were aligned to the same space, following the same pipeline for autofluorescence template reconstruction. Labeled neuron distributions were directly assigned to M132 and SARM parcellated areas within the CynoBrain reference space. To quantitatively evaluate both intrahemispheric and interhemispheric connectivity, heatmaps of logarithmic FLN (Fraction of Labeled Neurons) and volume-normalized projection density were generated across ipsilateral and contralateral parcellated areas.

**Figure 5.**
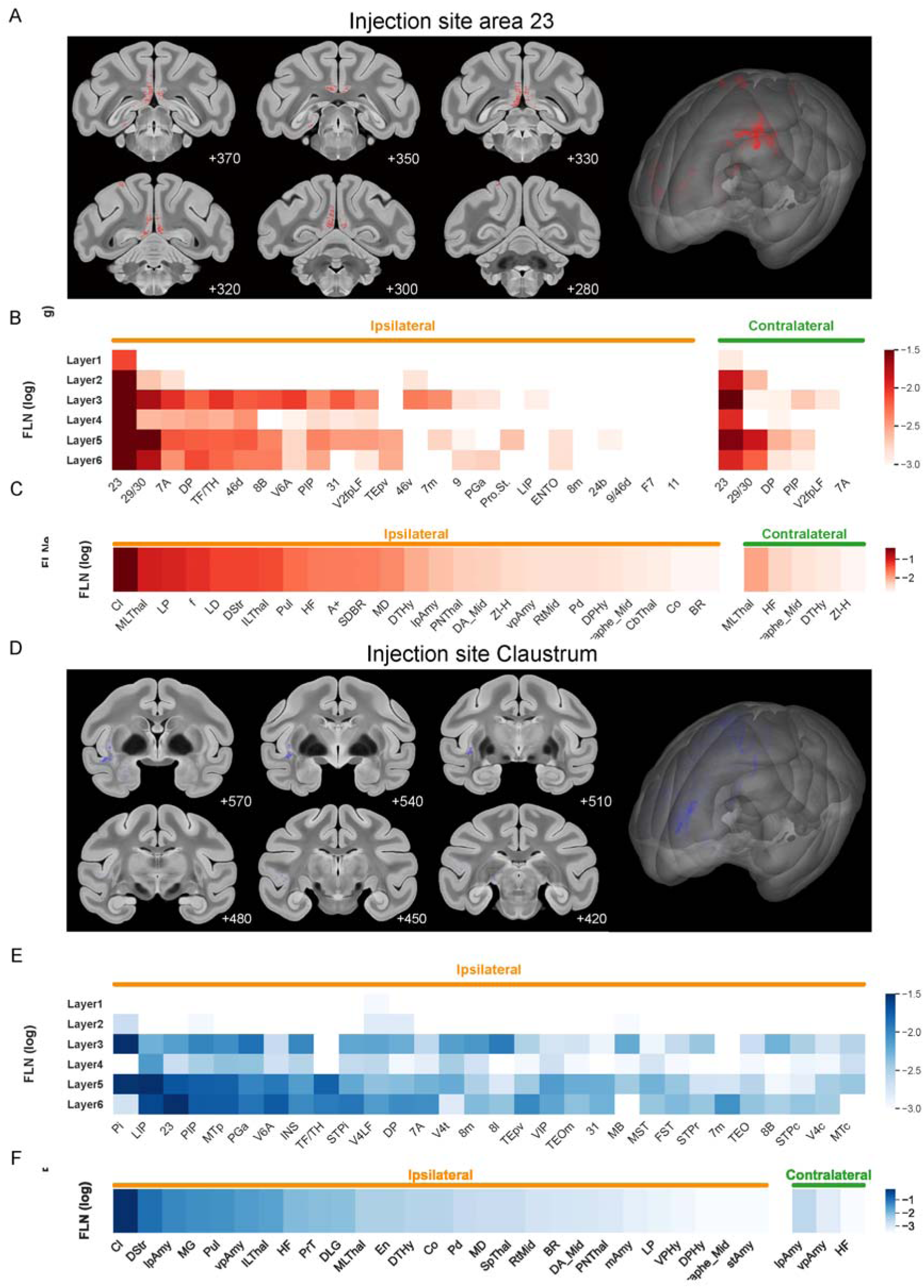
Spatial distribution of retrograde tracing data reconstructed to CynoBrain space. **A** Reconstructed coronal sections (+280 to +370) showing the distribution of labeled neurons by tracer injection in area 23 of the posterior cingulate cortex (red), with a 3D cortical surface reconstruction illustrating the distribution of labeled neurons. **B-C** Heatmaps showing the FLN of retrogradely labeled neurons (log scale) across ipsilateral and contralateral cortical areas (B) and subcortical areas (C) projecting to area 23. **D** Reconstructed coronal sections (+420 to +570) showing the distribution of labeled neurons by tracer injection in the claustrum (blue), with a 3D surface reconstruction of labeled neurons distribution. **E-F** Heatmaps showing the projection density (E) and FLN (F) of retrogradely labeled neurons (log scale) across ipsilateral and contralateral cortical areas projecting to claustrum. Cortical parcellation is based on the M132 105-area parcellation and subcortical parcellation is based on the SARM.

For injection site aera 23, labeled neurons were identified across coronal sections spanning from the level of the anterior commissure to the genu of the corpus callosum (Fig. 5A and Supplementary Fig. 4). Three-dimensional surface rendering of the labeled neuron distributions was concentrated along the medial cortical wall (Fig. 5A, Right). Cortical input analysis revealed widespread projections to area 23 from multiple ipsilateral cortical regions, with the strongest FLN values observed in cingulate, medial prefrontal, temporal, and posterior parietal association areas (Fig. 5B and Supplementary Fig. 5A). Detailed values were listed in Supplementary Table 3, 4 and 5. Layer-resolved heatmaps showed that these cortical inputs were distributed across multiple cortical layers, with stronger labeling generally observed in Layers 3, 5, and 6. The highest ipsilateral cortical contributions were detected from area 23, area 29/30, 7A, DP, TF/TH, 46d, 8B, V6A, PIP, and area 31. Contralateral cortical projections were also evident but were more restricted and generally weaker than ipsilateral inputs, mainly originating from homologous or functionally related regions including area 23, area 29/30, DP, PIP, and V2fpLF.

Subcortical projections to area 23 were also detected across a range of ipsilateral and contralateral structures (Fig. 5C and Supplementary Fig. 6). Ipsilateral subcortical inputs showed prominent FLN values in thalamic, striatal, limbic, and midbrain-related regions, including MLThal, LP, LD, H.Thal, Put, HF, SDBR, MD, DTHy, lpAmy, PNThal, DA_Mid, ZI-H, vpAmy, RtMid, Pd, DPHy, aphe_Mid, CbThal, Co, and BR. Compared with the cortical projection pattern, contralateral subcortical labeling was more limited, with detectable signals mainly observed in MLThal, HF, aphe_Mid, DTHy, and ZI-H. These results indicate that area 23 receives broad ipsilateral cortical and subcortical inputs, accompanied by weaker but distinguishable contralateral projections.

For the claustrum injection site, retrogradely labeled neurons were registered to CynoBrain space across coronal sections spanning +420 to +570 (Fig. 5D). Labeled neurons were distributed primarily in lateral cortical and insular territories, and the 3D surface reconstruction showed a lateralized distribution pattern distinct from the medial distribution observed for the area 23 injection. Cortical projection analysis demonstrated that claustrum-projecting neurons were broadly distributed across ipsilateral cortical areas (Fig. 5E and Supplementary Fig. 5B). Detailed values were listed in Supplementary Table 6, 7 and 8. Layer-resolved FLN heatmaps showed prominent labeling in Layers 3, 5, and 6, suggesting that claustral inputs arise from both supragranular and infragranular cortical layers. The strongest ipsilateral cortical projections originated from parietal, temporal, insular, and visual association cortices, including LIP, PI, 23, PIP, MTn, PGa, V6A, INS, TF/TH, STPi, VAL, DP, 7A, V4t, 8m, TEpv, VIP, TEOm, 31, MB, MST, FST, STPr, 7m, TEO, 8B, STPc, V4c, and MTc. In contrast to area 23, contralateral cortical projections to the claustrum were weak or not apparent in the cortical heatmap, indicating a strong ipsilateral dominance of cortical inputs.

Subcortical input analysis further revealed extensive ipsilateral projections to the claustrum from multiple subcortical systems (Fig. 5F). Strong FLN values were observed in CI, DStr, lpAmy, MG, Pul, vpAmy, ILThal, HF, PPT, DLG, MLThal, En, DTHy, Co, Pd, MD, SpThal, RtMid, BR, DA_Mid, PNThal, mAmy, LP, VPHy, DPHy, aphe_Mid, and stAmy. These inputs included thalamic nuclei, basal ganglia-related regions, amygdalar nuclei, hippocampal formation, and midbrain-related structures, indicating broad subcortical convergence onto the claustrum. A small contralateral subcortical component was also detected, primarily from lpAmy, vpAmy, and HF, but these inputs were substantially weaker than ipsilateral projections. Together, these findings demonstrate distinct organizational patterns between the two injection sites: area 23 receives bilateral cortical and subcortical inputs with an ipsilateral bias, whereas the claustrum receives predominantly ipsilateral cortical and subcortical inputs with only limited contralateral contribution. Brain region abbreviations used are provided with their full names in Supplementary Tables 9–11.

Collectively, these findings establish that our unified CynoBrain framework provides a robust and standardized platform for whole-brain atlasing of retrograde tracing data.

### 2.5. An Interactive Online Platform of CynoBrain Atlas

To make the full suite of CynoBrain datasets, we developed the Digital Brain CynoBrain online platform (https://macaque.digital-brain.cn/cynobrain-atlas/). This platform was designed to visualize the comprehensive anatomical and parcellation datasets generated in the present study, organized according to multiple hierarchical parcellation schemes, including the D99, M132, and SARM atlases, and is hosted on a cloud-based service-oriented architecture.

The CynoBrain platform allows users to interactively explore the macaque brain atlas across multiple levels of anatomical detail. In the 2D Viewer, users can browse the full set of coronal sections spanning the anterior-to-posterior extent of the brain, with parcellation labels overlaid on the cortical parcellations. In the 3D Viewer, users can select individual brain regions or entire lobes from a hierarchically organized region list, encompassing frontal, parietal, and other major cortical divisions along with their constituent subregions and render interactive three-dimensional surface models of the selected structures overlaid on the template (Fig. 6). Users can further adjust the transparency, color, and visibility of individual regions, and query region-specific metadata including anatomical abbreviations and full nomenclature.

**Figure 6.**
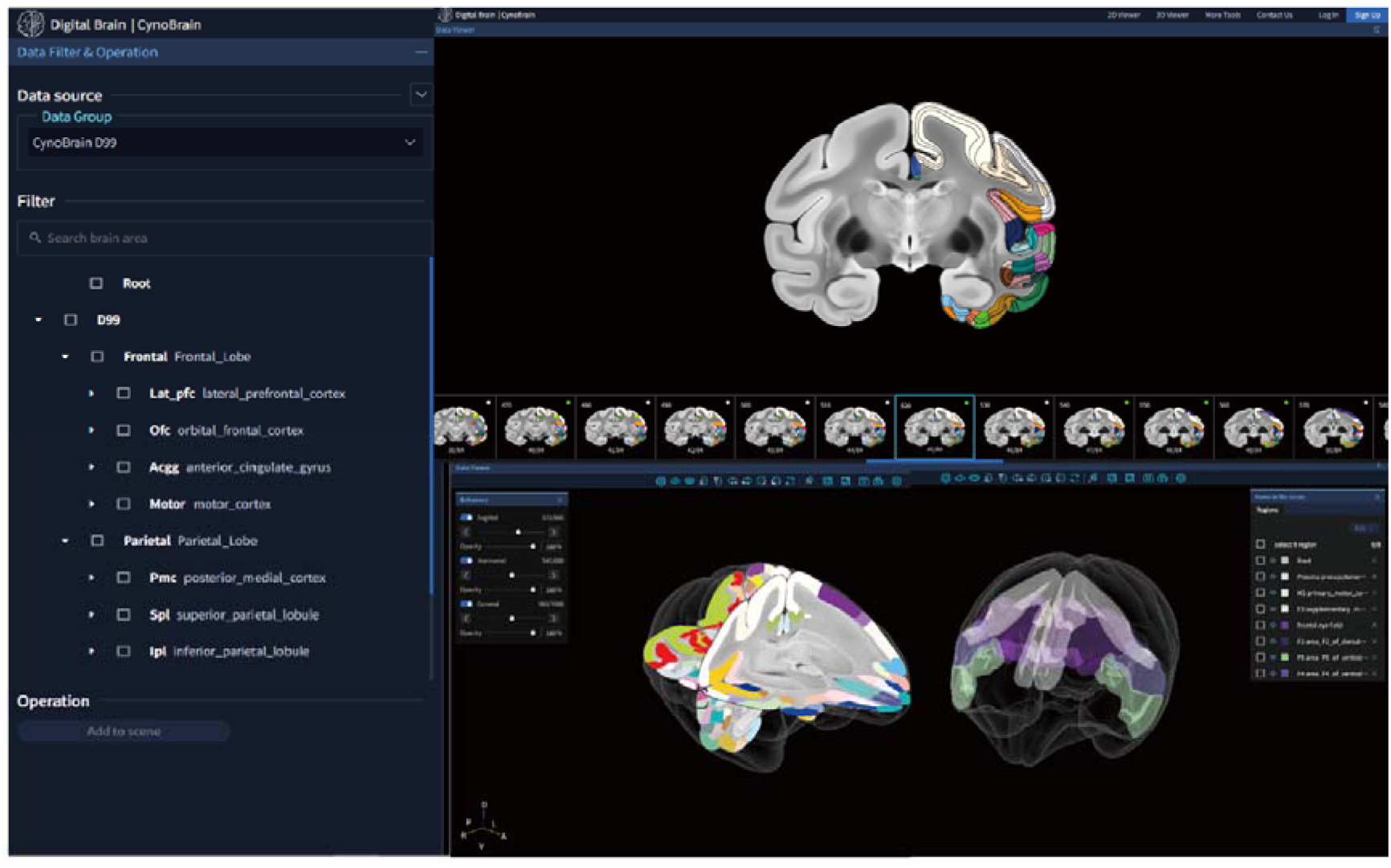
Interactive online platform of CynoBrain Atlas. Screenshot of the web-based CynoBrain platform showing the hierarchical region list and anatomical parcellations in different views. The platform enables interactive exploration of the macaque brain atlas through 2D coronal sections and 3D surface renderings. The viewer allows precise visualization of selected brain structures on the whole-brain template, and provides tools to adjust region transparency, color, and visibility while querying anatomical metadata.

## 3. Discussion

In the present study, we developed CynoBrain, a high-resolution multimodal brain atlas and an integrated platform capable of anchoring diverse data types within a common anatomical space. The atlas combines a 75 µm isotropic MRI template, a complementary autofluorescence histological template, multi-atlas cortical and subcortical parcellations, transcriptomics-derived laminar definition, and corticocortical connectivity data. This integrative architecture addresses a fundamental limitation of existing macaque atlases, which have largely been developed as modality-specific resources[21, 29, 30] and have not provided a standardized infrastructure for cross-modal data fusion[31].

The resolving power of the CynoBrain high-resolution template represents a substantial advance over previous macaque reference spaces. Existing templates, including the INIA19, Yerkes19 and NMT v2, were constructed at resolutions ranging from 150 to 500 µm isotropic, which are insufficient to resolve thin laminar structures or reliably delineate boundaries of small subcortical nuclei. In contrast, the high resolution of CynoBrain template (at 75µm isotropic) allows delineation of the reticular nucleus of the thalamus [32], the visualization of the claustrum as a discrete gray matter sheet separated from adjacent white matter [33], and the identification of the Stria of Gennari within V1 as a hyperintense stripe within the cortical ribbon [34]. The visibility of the Stria of Gennari is particularly significant: it enables direct MRI-based delineation of V1 boundaries without recourse to histological sections, a result with direct implications for the MRI-based parcellation of early visual cortex[35, 36].

The accompanying autofluorescence template of CynoBrain is also a key advance over current MRI-based macaque atlases, as none of them incorporate a histological modality as a co-registered reference [21, 29, 30]. The clear delineation of hippocampal subfields CA1, CA2, and CA3[37, 38], and the laminar structure of the lateral geniculate nucleus, in the autofluorescence template clearly demonstrate a complementary contrast to MRI. By registering both modalities into a common reference space, CynoBrain enables researchers to leverage MRI for volumetric coverage and geometric standardization while drawing on autofluorescence imaging to resolve cytoarchitectural boundaries where MRI contrast is insufficient.

The multi-atlas parcellation incorporated in CynoBrain framework reflects the practical advantage for a single reference space. By making three major parcellation schemes simultaneously available within a single template space, the CynoBrain framework enables direct comparison of parcellation-dependent results across studies that have adopted different areal definitions. Of particular significance is the volumetric reconstruction of the M132 105-area cortical parcellation. This parcellation has been primarily distributed as a surface-based representation, and no standardized volumetric version was available before. The surface-informed label optimization procedure employed here ensures that the volumetric projection faithfully preserves the areal boundaries defined by the original cytoarchitectonic criteria. The M132 scheme serves as the foundation of the Kennedy–Markov quantitative connectivity dataset, which provides the most comprehensive quantitative directed connectivity matrix available for any primate brain [6, 39]. However, despite its unparalleled comprehensiveness, integrating this dataset with in vivo neuroimaging has remained methodologically inconvenient: without a standardized volumetric representation of M132, individual studies have relied on ad hoc surface-to-volume projections that vary in methodology, undermining the reproducibility and cross-study comparability that the dataset’s quantitative precision is intended to support. The volumetric M132 available through CynoBrain enables direct quantitative comparison between tracer-derived connectivity weights and in vivo measures of structural or functional connectivity within the same parcellation framework [40, 41], and finally bridges the gap between a wealth of historical retrograde/anterograde tracing data and modern in vivo neuroimaging modalities within a single reference frame [15, 42–44].

The integration of transcriptomics-defined laminar maps provides a new feature that is not available in other macaque atlases. Existing macaque atlases have not defined cortical layers systemically, and thus current macaque brain mapping datasets can only provide spatial information at region level. However, laminar organization is no doubt a fundamental and critical features of primate cortex, and spatial information of macaque brain mapping datasets would be significantly enriched if laminar definition is incorporated as in the current CynoBrain platform. Importantly, such laminar definition is largely data-driven based on the macaque whole-brain spatial transcriptomic data [12], providing a unbiased cortical layering scheme.

The multimodal integration capability of the CynoBrain platform is clearly demonstrated by mapping retrograde tracing data into this reference space. Conventionally, current macaque connectivity databases, including CoCoMac[15], aggregate findings at brain region level, but do not provide spatially registered distributions within a common volumetric reference space [45]. By transforming labeled neuron coordinates into CynoBrain template space using the current 3D reconstruction pipeline, the present approach provides the capability of constructing a unified anatomical connectivity database, similar to that in the mouse brain [46]. Such parcellation independent connectivity database in a common reference space would be greatly beneficial to the primate neuroscience community, as much richer information can be further extracted by the whole community, compared to a simple connectivity matrix. Furthermore, the current CynoBrain platform also provide convenient integration with other brain mapping datasets, for example structural, functional and diffusion MRI[47].

Looking forward, the CynoBrain framework is designed as an open and expandable platform rather than a static atlas resource. The standardized registration workflow and cloud-based online platform provide the infrastructure necessary to incorporate additional data types as they become available — including resting-state and task-based functional MRI connectivity maps[31, 48], diffusion MRI tractography[40, 49], single-cell and spatial transcriptomic profiles[50, 51], and electrophysiological recordings, all progressively building toward a comprehensive, multimodal macaque brain data platform. Initiatives such as the China Brain Project are generating large-scale macaque datasets across multiple modalities and laboratories. The CynoBrain framework provides a natural integration platform for these efforts, enabling cross-study normalization and cumulative knowledge synthesis within a species-matched, high-resolution reference space.

## 4. Methods

### 4.1. Animals and Ethical Statement

All animal experiments were approved by the Animal Care and Use Committee of the Institute of Neuroscience, Chinese Academy of Sciences, Shanghai, China, and conducted in accordance with institutional guidelines. A total of 22 adult cynomolgus macaques (Supplementary Table 1) were used across the studies described in this work. The sample was predominantly female (n = 21, one male), with ages ranging from 7 to 16 years (mean = 11.23 ± 1.85 years) and weight from 2.9 to 7.1kg (mean = 4.60 ± 1.23 kg). Efforts were made to minimize animal suffering, and animal care was supervised by technicians skilled in primate healthcare and housing. Specimens were obtained at necropsy immediately following euthanasia performed for reasons unrelated to the present study. Following extraction from the skull, brain specimens were transferred to phosphate-buffered saline (PBS).

A subset of 13 animals (one male) was chosen for autofluorescence template construction. These animals were premedicated with atropine (0.05 mg/kg) and dexamethasone (0.5 mg/kg), and anesthesia was induced with Zoletil (Virbac, 5 mg/kg) and maintained with 1.5–2.5% isoflurane in pure oxygen. Retrograde tracers, FastBlue (Polysciences, 2–3% in water), cholera toxin subunit B conjugated with Alexa Fluor (Molecular Probes, 1–3% in PBS), and Diamidino Yellow (2–3% in water), were injected into targeted brain regions under guidance of structural MRI (Siemens 3T Prisma). An image-based neuronavigation system (Brainsight, Rogue Research Inc.) was employed to facilitate accurate targeting of deep brain structures. Following a survival period of 11–13 days, animals were deeply anesthetized and transcardially perfused with saline, 4% paraformaldehyde (PFA), and PB-glycerine solution (5%).

### 4.2. Construction of the CynoBrain MRI Template

#### 4.2.1. MRI Data Acquisition

Prior to imaging, brain specimens were submerged in liquid fluorocarbon (Galden PFPE, Solvay, Brussels, Belgium) and placed under vacuum conditions for approximately 12 hours to remove residual air bubbles trapped within the cortical sulci, thereby minimizing susceptibility artifacts during MRI acquisition. All MRI acquisitions were performed on a Bruker 9.4 Tesla scanner (Bruker Bio Spin, USA). Images were acquired using a 3D FLASH pulse sequence with the following parameters: TR/TE = 100/13 ms, flip angle = 20°, averages = 4, matrix = 440 × 380 × 300, FOV = 66.0 × 57.0 × 45.0 mm³, resolution = 0.15 × 0.15 × 0.15 mm³, coronal orientation with Ro-Cd readout direction, with a total scanning time of approximately 12 hours 40 minutes per specimen.

#### 4.2.2. Generation of the Symmetric CynoBrain Template

The CynoBrain template was constructed following a symmetric iterative registration pipeline adapted from the NMT v2 workflow[52]. Prior to registration, each individual anatomical scan was upsampled by a factor of 2 to an isotropic voxel resolution of 75 µm using sinc interpolation. To enforce bilateral symmetry in the final template, each upsampled volume was duplicated and mirrored about the interhemispheric plane, with both the original and mirrored copies treated as independent inputs to the registration pipeline.

Template construction proceeded through 5 terative cycles of groupwise nonlinear registration. In each iteration, all input volumes were first affinely aligned to correct for global differences in position, orientation, and scale, and then nonlinearly registered to the current intermediate template using ANTs[53, 54], employing a symmetric diffeomorphic normalization (SyN) algorithm with Mattes mutual information as the similarity metric. Brain masking was applied throughout to restrict registration to brain tissue. The registered volumes were averaged to produce an updated template for the subsequent iteration. This process progressively sharpened anatomical contrast and reduced individual-specific bias, converging toward a population-representative mean anatomy. The resulting CynoBrain template has a final isotropic voxel resolution of 75 µm, and the enforced left-right symmetry ensures the template is free of lateralization biases, making it suitable as a neutral reference space for group-level normalization and atlas alignment.

#### 4.2.3. Alignment of Cortical and Subcortical Parcellations

Two established parcellations, D99 atlas and SARM parcellation, were transferred from NMT v2 space into CynoBrain template space via nonlinear deformable registration using ANTs (SyN algorithm), followed by slice-by-slice manual refinement to correct residual misalignments. For D99 atlas, parcellation was restricted to the 143 cortical areas defined in Chen et al. [12].

The M132 105-area parcellation, originally defined only on 2D cortical images, was additionally projected into the CynoBrain volume. To ensure parcellation boundaries were oriented perpendicular to the cortical surface and consistent with the columnar organization of the cerebral cortex, a label optimization procedure was applied. Cortical depth was estimated by solving the Laplace equation within the gray matter volume, yielding a smooth radial orientation field across the cortical ribbon. Streamlines were then traced from the white matter surface toward the pial surface, and each voxel was assigned the dominant label of its corresponding streamline by majority vote, with isolated fragments reassigned to the most frequent neighboring label.

### 4.3. Reconstruction of retrograde tracing data to CynoBrain Space

#### 4.3.1. Data Acquisition

Following perfusion, brain tissues were dehydrated through a graded series of PB-glycerine solutions (10%, 15%, and 20%). Coronal sections (40 µm thickness) were obtained using the frozen section method, with every sixth section (240 µm interval) mounted on glass slides for connectivity analysis. Prior to each sectioning step, blockface images of the tissue block were acquired to provide a continuous photographic reference for subsequent 3D spatial reconstruction. Sections were digitized using an Axio Scan.Z1 slide scanner (Zeiss) with a 10× objective. Fluorescence filter settings were as follows: 670–710 nm (CTB647), 582–640 nm (CTB555), 507–543 nm (CTB488), and 430–470 nm (FastBlue). Retrogradely labeled neurons were identified and plotted using an in-house developed software package, CellPlot.

#### 4.3.2. Volumetric Reconstruction and Spatial Normalization

Histological sections, originally digitized at 0.65 µm resolution, were downsampled to 25 µm for 3D reconstruction, with a section interval of 240 µm. Prior to registration, all sections underwent correction for illumination imbalances and optical artifacts to ensure consistent image quality across the sections.

The 3D shape of the postmortem brain was reconstructed through a hierarchical series of registration steps using ANTs. First, each histological section was nonlinearly registered to its corresponding blockface image on a section-by-section basis using ANTs (SyN algorithm), compensating for local tissue distortions introduced during cryosectioning and mounting. The corrected sections were then assembled into a coarse 3D volume, which was subsequently aligned to the postmortem MRI dataset via 3D affine registration using ANTs to establish a common anatomical reference frame. Following this global alignment, each section was further refined through 2D nonlinear registration to its corresponding MRI slice using ANTs (SyN algorithm), correcting residual local deformations and substantially improving the overall fidelity of the reconstruction.

Upon completion of reconstruction, the dataset was spatially normalized to the CynoBrain template via 3D deformable registration using ANTs (SyN algorithm). The resulting deformation fields were stored for each reconstructed dataset. The spatial coordinates of retrogradely labeled neurons plotted with CellPlot were subsequently transformed into CynoBrain template space by applying the corresponding stored transformation fields, enabling direct cross-individual comparison of projection neuron distributions.

#### 4.3.3. Quantification of Projection Connectivity

To visualize projection patterns across the cortical regions, connectivity heatmaps were generated for each injection case in CynoBrain template space. The transformed coordinates of retrogradely labeled neurons were aggregated within each parcellated area defined by the CynoBrain atlas. The connectivity metrics were computed for each source region: (1) the fraction of labeled neurons (FLN), defined as the proportion of labeled neurons in a given source area relative to the total number of projections.

FLN values were logarithmically transformed (log) prior to visualization to accommodate their wide dynamic range. Heatmaps were constructed separately for ipsilateral and contralateral projections, with parcellated areas and injection cases arranged along the two axes. Color intensity represented either log (FLN) or volume-normalized projection density using a perceptually uniform colormap. Areas with no detected labeled neurons were assigned a floor value to distinguish true absence from weak connectivity.

### 4.4. Laminar Delineation of the Macaque Cortex

#### 4.4.1. Transcriptomic–MRI Co-registration

A computational framework was developed to integrate layer-specific gene expression data[12] with the CynoBrain high-resolution MRI template. The pipeline combined slice-to-volume registration, which established spatial correspondence between transcriptomic sections and the MRI volume, with surface-based laminar delineation to enable precise mapping of cortical layers across the entire cortex.

The registration pipeline proceeded in four stages: identification of the best-matching MRI reference slices based on structural similarity metrics, rigid registration to preserve overall anatomical shape, affine registration to account for scaling and shearing differences, and finally nonlinear deformable registration using ANTs (SyN algorithm) to correct for local tissue distortions. Sections exhibiting residual misalignment after automated registration were identified through visual inspection and corrected via manual landmark-based adjustment, ensuring anatomical accuracy across all sections.

#### 4.4.2. Surface-Based Laminar Delineation

Laminar delineation began with surface extraction from the MRI volume to generate an initial cortical mesh. The mesh underwent iterative refinement to optimize surface accuracy and minimize geometric artifacts, followed by Laplacian smoothing to reduce noise while preserving anatomical features. The processed surface was then projected back onto the volumetric space, enabling the generation of a comprehensive 3D laminar delineation map spanning the entire macaque cortex.

#### 4.4.3. Measurement of layer-specific cortical thickness

Layer-specific thickness measurement. Cortical layer thickness was quantified from 2D sections across 143 regions. For each region, layer-specific thickness values were extracted from pixels within the region mask. To remove measurement artifacts while preserving genuine thin cortex, we applied a two-tier outlier removal strategy targeting. At the slice level, we calculated median thickness and median absolute deviation (MAD) for each layer. Slices exceeding M+2.0×MAD were excluded. At the pixel level, remaining data were filtered using M+2.0×MAD threshold. Final layer thickness was computed as the mean of retained pixels, and total thickness as the sum of six layers.

## Acknowledgements

The project was supported by National Science and Technology Innovation 2030 Major Program (2021ZD0200100), Shanghai Municipal Science and Technology Major Project (2018SHZDZX05), National Key R&D Program of China (2022YEF0203200), Strategic Priority Research Program of CAS (XDB32010100 and XDA27010400), Lingang Laboratory (LG202105-01-01, LG202104-02, LG202104-02-02, LG202104-02-03, LG202104-02-04, and LG202104-02-06), Shanghai Municipal of Science and Technology Project (21015800100), International Partnership Program of Chinese Academy of Sciences (153D31KYSB20170059), Scientific Instrument Developing Project of CAS (YJKYYQ20190052), Scientific Instrument Developing Project of NNSFC (31827803), National Natural Science Foundation of China (31900466 and 32161133024), and Shenzhen Basic Research Project for Excellent Young Scholars (2020251518).

## Author Contributions

Z.L. and Z.S. designed and supervised the study; M.W. N.Y. performed the animal surgery; H.Y., M.P., B.B., K.Z. and X.Y collected the MRI data; H.Y., M.W. and Z.L. conducted the data analysis. Y.H., M.W. and Z.M. verified the underlying data and wrote the manuscript. All authors discussed and commented on the manuscript and approved the final version for submission.

## Data Availability Statement

CynoBrain is openly accessible (https://macaque.digital-brain.cn/cynobrain-atlas/). All data collected for the study, including raw data and code for data analysis will be made available upon reasonable request.

