## Supplementary file for "CynoBrain: A unified high-resolution framework for cross-modal data integration for macaque brain mapping"

Title

**Table S1. Information of macaque subjects**

| **Monkey ID** | **Age (year)** | **Weight (kg)** | **Sex** |
| --- | --- | --- | --- |
| ***C056*** | 10 | 4.1 | Female |
| ***C057*** | 10 | 5.2 | Female |
| ***C058*** | 10 | 3.6 | Female |
| ***C059*** | 10 | 4.0 | Female |
| ***C060*** | 7 | 5.6 | Male |
| C063 | 12 | 4.8 | Female |
| ***C064*** | 10 | 3.2 | Female |
| ***C067*** | 14 | 3.6 | Female |
| ***C068*** | 12 | 7.1 | Female |
| C069 | 12 | 3.8 | Female |
| ***C070*** | 11 | 3.3 | Female |
| ***C071*** | 16 | 3.5 | Female |
| ***C072*** | 11 | 3.4 | Female |
| ***C073*** | 11 | 4.0 | Female |
| ***C074*** | 10 | 4.6 | Female |
| C075# | 10 | 4.4 | Female |
| C088 | 11 | 2.9 | Female |
| C091 | 13 | 4.3 | Female |
| C107 | 13 | 6.5 | Female |
| C108 | 13 | 6.0 | Female |
| C109 | 10 | 5.2 | Female |
| C111 | 11 | 5.8 | Female |
| C112 | 10 | 6.6 | Female |

*Bold italic indicates subjects included in the autofluorescence template group. Macaque C075 was not used for template rescontruction.

**Table S2. Cortical thickness measurements across 143 cortical regions**

| **Region** | **Layer I (mm)** | **Layer II (mm)** | **Layer III (mm)** | **Layer IV (mm)** | **Layer V (mm)** | **Layer VI (mm)** | **Total (mm)** |
| --- | --- | --- | --- | --- | --- | --- | --- |
| 1-2 | 0.157117 | 0.160785 | 0.558291 | 0.155508 | 0.325551 | 0.480373 | 1.837625 |
| 10mc | 0.24763 | 0.19264 | 0.683251 | 0.098282 | 0.326936 | 0.361925 | 1.910664 |
| 10mr | 0.211744 | 0.178378 | 0.780156 | 0.131892 | 0.361837 | 0.577639 | 2.241645 |
| 10o | 0.202319 | 0.169719 | 0.63466 | 0.145665 | 0.311165 | 0.465543 | 1.929071 |
| 11l | 0.202622 | 0.166593 | 0.655088 | 0.117095 | 0.276061 | 0.40256 | 1.82002 |
| 11m | 0.207221 | 0.176798 | 0.640789 | 0.114233 | 0.236554 | 0.307306 | 1.6829 |
| 12l | 0.289896 | 0.180917 | 0.899061 | 0.200106 | 0.417574 | 1.157303 | 3.144855 |
| 12m | 0.253678 | 0.163333 | 0.931041 | 0.188092 | 0.370589 | 0.966613 | 2.873346 |
| 12o | 0.340405 | 0.209563 | 0.903891 | 0.16247 | 0.483924 | 0.890751 | 2.991005 |
| 12r | 0.231018 | 0.187718 | 0.764041 | 0.158093 | 0.339159 | 0.738685 | 2.418714 |
| 13a | 0.312853 | 0.182085 | 0.517578 | 0 | 0.503209 | 0.481669 | 1.997395 |
| 13b | 0.203704 | 0.164445 | 0.606498 | 0 | 0.366041 | 0.422306 | 1.762994 |
| 13l | 0.490827 | 0.22337 | 1.119507 | 0 | 0.504572 | 0.86119 | 3.199467 |
| 13m | 0.264304 | 0.214909 | 0.909304 | 0 | 0.539431 | 0.88101 | 2.808957 |
| 14c | 0.190471 | 0.153042 | 0.387411 | 0 | 0.431233 | 0.506692 | 1.668849 |
| 14r | 0.200798 | 0.150626 | 0.462822 | 0 | 0.294663 | 0.542277 | 1.651185 |
| 23a | 0.180394 | 0.187976 | 0.511332 | 0.13569 | 0.403302 | 0.520451 | 1.939146 |
| 23b | 0.176789 | 0.168565 | 0.563223 | 0.130217 | 0.396445 | 0.496069 | 1.931307 |
| 23c | 0.151301 | 0.170032 | 0.535024 | 0.10863 | 0.280488 | 0.270187 | 1.515661 |
| 24a | 0.194227 | 0.185868 | 0.403387 | 0 | 0.346939 | 0.296443 | 1.426865 |
| 24a' | 0.200299 | 0.157257 | 0.482664 | 0 | 0.367794 | 0.392492 | 1.600507 |
| 24b | 0.188887 | 0.187502 | 0.523861 | 0 | 0.592519 | 0.754873 | 2.247642 |
| 24b' | 0.180543 | 0.156227 | 0.490066 | 0 | 0.538733 | 0.77749 | 2.143058 |
| 24c | 0.168495 | 0.190316 | 0.523642 | 0 | 0.381345 | 0.336481 | 1.600279 |
| 24c' | 0.134364 | 0.198933 | 0.59923 | 0 | 0.417376 | 0.413926 | 1.763828 |
| 25 | 0.294502 | 0.164376 | 0.484189 | 0 | 0.352983 | 0.31024 | 1.606289 |
| 29 | 0.175888 | 0.344339 | 0.284225 | 0 | 0.537199 | 0.280957 | 1.622607 |
| 30 | 0.210079 | 0.199365 | 0.60761 | 0.106368 | 0.480415 | 0.542379 | 2.146215 |
| 31 | 0.167532 | 0.177836 | 0.682972 | 0.144898 | 0.357004 | 0.416168 | 1.94641 |
| 32 | 0.216504 | 0.18173 | 0.778534 | 0 | 0.509124 | 0.556092 | 2.241986 |
| 35 | 0.21346 | 0.190683 | 0.649212 | 0 | 0.448138 | 0.793365 | 2.294857 |
| 36c | 0.192436 | 0.175619 | 0.721767 | 0.128005 | 0.441167 | 0.770776 | 2.429771 |
| 36p | 0.210485 | 0.229264 | 0.84815 | 0.167113 | 0.650934 | 0.955857 | 3.061803 |
| 36r | 0.222284 | 0.182759 | 0.678711 | 0.167059 | 0.628096 | 1.620375 | 3.499285 |
| 3a-b | 0.155296 | 0.174353 | 0.595003 | 0.159096 | 0.252161 | 0.367258 | 1.703167 |
| 44 | 0.351227 | 0.169441 | 0.645133 | 0 | 0.265357 | 0.303197 | 1.734355 |
| 45a | 0.230637 | 0.185213 | 0.81318 | 0.189278 | 0.465113 | 1.084414 | 2.967835 |
| 45b | 0.179975 | 0.180256 | 0.810456 | 0.157617 | 0.429955 | 0.812748 | 2.571008 |
| 46d | 0.175648 | 0.169913 | 0.663485 | 0.126847 | 0.283027 | 0.402664 | 1.821583 |
| 46v | 0.168001 | 0.176895 | 0.640754 | 0.130205 | 0.263467 | 0.332721 | 1.712043 |
| 7op | 0.169009 | 0.192874 | 0.750611 | 0.153791 | 0.381094 | 0.477429 | 2.124807 |
| 8Ad | 0.152656 | 0.181756 | 0.667446 | 0.149786 | 0.366759 | 0.812066 | 2.330469 |
| 8Av | 0.157033 | 0.152362 | 0.555187 | 0.170719 | 0.370019 | 0.941112 | 2.346432 |
| 8Bd | 0.21314 | 0.19506 | 0.927722 | 0 | 0.621702 | 1.022911 | 2.980536 |
| 8Bm | 0.19999 | 0.190548 | 0.636497 | 0 | 0.57059 | 0.912764 | 2.51039 |
| 8Bs | 0.155533 | 0.192929 | 0.680685 | 0 | 0.347434 | 0.366618 | 1.743199 |
| 9d | 0.244516 | 0.180367 | 1.029591 | 0.130728 | 0.575525 | 0.964994 | 3.125722 |
| 9m | 0.205939 | 0.168308 | 0.807409 | 0.126321 | 0.464635 | 1.157659 | 2.930271 |
| AI | 0.160601 | 0.178114 | 0.660508 | 0.129275 | 0.317007 | 0.374795 | 1.820302 |
| AIP | 0.151176 | 0.187936 | 0.659657 | 0.11715 | 0.248015 | 0.251353 | 1.615288 |
| AL | 0.138615 | 0.16848 | 0.589412 | 0.153091 | 0.406346 | 0.646409 | 2.102353 |
| CL | 0.162848 | 0.194738 | 0.724456 | 0.159262 | 0.328909 | 0.477776 | 2.047989 |
| CM | 0.190311 | 0.189772 | 0.64303 | 0.113867 | 0.206197 | 0.265687 | 1.608864 |
| CPB | 0.177211 | 0.160069 | 0.707538 | 0.171423 | 0.423423 | 0.738218 | 2.377882 |
| DP | 0.198967 | 0.244132 | 0.724132 | 0.217964 | 0.434974 | 0.553896 | 2.374065 |
| EC | 0.303176 | 0.182948 | 0.756219 | 0 | 0.440551 | 0.727735 | 2.410629 |
| ECL | 0.367826 | 0.197979 | 0.884757 | 0 | 0.382635 | 0.703148 | 2.536346 |
| EI | 0.265773 | 0.160856 | 0.711614 | 0 | 0.383281 | 0.639277 | 2.1608 |
| ELc | 0.217746 | 0.183537 | 0.593914 | 0 | 0.376276 | 0.568794 | 1.940268 |
| ELr | 0.28278 | 0.203463 | 0.470013 | 0 | 0.304572 | 0.866729 | 2.127558 |
| EO | 0.198847 | 0.190066 | 0.707395 | 0 | 0.508913 | 0.525427 | 2.130648 |
| ER | 0.240371 | 0.213236 | 0.83723 | 0 | 0.649521 | 0.817838 | 2.758196 |
| F1 | 0.165065 | 0.178578 | 0.871788 | 0 | 0.81987 | 1.077672 | 3.112974 |
| F2 | 0.170811 | 0.201052 | 0.823365 | 0 | 0.73552 | 1.065968 | 2.996717 |
| F3 | 0.165568 | 0.181452 | 0.685255 | 0 | 0.64286 | 1.092648 | 2.767783 |
| F4 | 0.213774 | 0.227378 | 0.828281 | 0 | 0.767633 | 1.150491 | 3.187557 |
| F5 | 0.224666 | 0.193802 | 0.939112 | 0 | 0.659409 | 0.846633 | 2.863622 |
| F6 | 0.184135 | 0.188549 | 0.680546 | 0 | 0.585526 | 1.075384 | 2.714141 |
| F7 | 0.185976 | 0.173209 | 0.831306 | 0 | 0.714614 | 1.202331 | 3.107436 |
| FST | 0.1654 | 0.187745 | 0.658394 | 0.153105 | 0.322076 | 0.289165 | 1.775886 |
| G | 0.289372 | 0.203937 | 0.625716 | 0.102199 | 0.234209 | 0.237667 | 1.6931 |
| IPa | 0.242114 | 0.228827 | 0.716237 | 0.133411 | 0.222138 | 0.278215 | 1.820942 |
| Ia | 0.231554 | 0.176657 | 0.470017 | 0 | 0.368935 | 0.581813 | 1.828977 |
| Iai | 0.363955 | 0.235467 | 1.449699 | 0 | 0.781295 | 1.000369 | 3.830784 |
| Ial | 0.228503 | 0.231584 | 0.770228 | 0 | 0.508957 | 0.832523 | 2.571795 |
| Iam | 0.329681 | 0.166063 | 0.570362 | 0 | 0.548004 | 0.881933 | 2.496043 |
| Iapl | 0.203395 | 0.222635 | 0.708942 | 0 | 0.449135 | 0.320244 | 1.904352 |
| Iapm | 0.269317 | 0.210581 | 0.599298 | 0 | 0.46092 | 0.572673 | 2.11279 |
| Id | 0.177656 | 0.193255 | 0.646316 | 0.132545 | 0.386054 | 0.516829 | 2.052655 |
| Ig | 0.156921 | 0.191554 | 0.554854 | 0.124733 | 0.26562 | 0.330542 | 1.624225 |
| LIPd | 0.169313 | 0.23509 | 0.833504 | 0.172735 | 0.361167 | 0.417995 | 2.189803 |
| LIPv | 0.146981 | 0.205595 | 0.708072 | 0.155234 | 0.292347 | 0.319549 | 1.827778 |
| LOP | 0.148277 | 0.195501 | 0.640168 | 0.15446 | 0.329864 | 0.387943 | 1.856214 |
| MIP | 0.147513 | 0.196918 | 0.6967 | 0.159997 | 0.272173 | 0.341566 | 1.814867 |
| ML | 0.208244 | 0.176721 | 0.87501 | 0.169389 | 0.463859 | 0.697089 | 2.590313 |
| MST | 0.173499 | 0.236223 | 0.773243 | 0.169327 | 0.303333 | 0.320579 | 1.976203 |
| MT | 0.216976 | 0.2492 | 0.990055 | 0.232163 | 0.460968 | 0.359655 | 2.509016 |
| Opt | 0.186731 | 0.228718 | 0.784617 | 0.181077 | 0.418243 | 0.506543 | 2.30593 |
| PE | 0.168891 | 0.181024 | 0.680069 | 0.176375 | 0.399007 | 0.591533 | 2.196899 |
| PEa | 0.150723 | 0.190996 | 0.791033 | 0.166907 | 0.404195 | 0.484692 | 2.188544 |
| PEc | 0.165513 | 0.178945 | 0.777212 | 0.201311 | 0.406005 | 0.619714 | 2.348699 |
| PEci | 0.147769 | 0.178747 | 0.516554 | 0.114451 | 0.223468 | 0.257378 | 1.438368 |
| PF | 0.159456 | 0.166087 | 0.684485 | 0.157936 | 0.411511 | 0.640498 | 2.219973 |
| PFG | 0.182464 | 0.174293 | 0.619896 | 0.165362 | 0.447383 | 0.802937 | 2.392335 |
| PG | 0.171345 | 0.192568 | 0.6475 | 0.192052 | 0.510133 | 0.741571 | 2.455168 |
| PGa | 0.234275 | 0.216492 | 0.714548 | 0.121168 | 0.219621 | 0.274405 | 1.780508 |
| PGm | 0.177268 | 0.19146 | 0.707709 | 0.146726 | 0.274871 | 0.385444 | 1.883479 |
| PIP | 0.170753 | 0.186109 | 0.50587 | 0.127412 | 0.181592 | 0.226942 | 1.398678 |
| Pi | 0.305631 | 0.174008 | 0.362172 | 0 | 0.206117 | 0.255834 | 1.303762 |
| Pir | 0.297473 | 0.294773 | 0.181069 | 0 | 0.218228 | 0.30556 | 1.297102 |
| PrCO | 0.199086 | 0.197293 | 0.825393 | 0 | 0.646138 | 0.736056 | 2.603966 |
| R | 0.170873 | 0.163488 | 0.546339 | 0.14717 | 0.304453 | 0.45216 | 1.784483 |
| RM | 0.263933 | 0.175901 | 0.561177 | 0.097765 | 0.205116 | 0.252178 | 1.55607 |
| RPB | 0.163094 | 0.154174 | 0.721108 | 0.158244 | 0.45615 | 0.690917 | 2.343688 |
| RT | 0.171376 | 0.191468 | 0.528111 | 0.145621 | 0.325975 | 0.515317 | 1.877868 |
| RTL | 0.165257 | 0.182429 | 0.620101 | 0.166749 | 0.469784 | 0.991572 | 2.595892 |
| RTM | 0.278 | 0.209273 | 0.648372 | 0.103289 | 0.273589 | 0.38475 | 1.897272 |
| RTp | 0.172509 | 0.207082 | 0.62505 | 0.176911 | 0.458634 | 0.776506 | 2.416693 |
| Ri | 0.327432 | 0.188181 | 0.576274 | 0.121302 | 0.187933 | 0.222135 | 1.623256 |
| SII | 0.142662 | 0.19274 | 0.654735 | 0.150223 | 0.306447 | 0.439065 | 1.885871 |
| STGr | 0.187302 | 0.184625 | 0.712753 | 0.139479 | 0.455811 | 0.689589 | 2.369559 |
| TAa | 0.17689 | 0.208206 | 0.81816 | 0.191316 | 0.481335 | 0.786087 | 2.661995 |
| TEO | 0.190866 | 0.206711 | 0.757066 | 0.18181 | 0.408104 | 0.445398 | 2.189955 |
| TEa | 0.182806 | 0.209679 | 0.831524 | 0.167052 | 0.356198 | 0.459807 | 2.207066 |
| TEad | 0.193621 | 0.178186 | 0.663098 | 0.172338 | 0.397088 | 0.649853 | 2.254183 |
| TEav | 0.214853 | 0.200272 | 0.777257 | 0.159827 | 0.467677 | 0.601519 | 2.421405 |
| TEm | 0.160141 | 0.202009 | 0.772878 | 0.18804 | 0.440796 | 0.619654 | 2.38352 |
| TEpd | 0.195754 | 0.170877 | 0.64771 | 0.15402 | 0.41972 | 0.520702 | 2.108783 |
| TEpv | 0.21959 | 0.17942 | 0.690646 | 0.140969 | 0.424886 | 0.48282 | 2.138331 |
| TF | 0.274722 | 0.189879 | 0.890469 | 0.139059 | 0.568704 | 0.736945 | 2.799778 |
| TFO | 0.249603 | 0.184961 | 0.678048 | 0.130213 | 0.336248 | 0.380524 | 1.959598 |
| TGa | 0.281295 | 0.256147 | 0.928564 | 0 | 0.69392 | 0.666942 | 2.826868 |
| TGdd | 0.245666 | 0.229859 | 0.774393 | 0 | 0.657852 | 0.553589 | 2.461359 |
| TGdg | 0.26557 | 0.252866 | 0.860685 | 0.148814 | 0.653301 | 0.477596 | 2.658831 |
| TGsts | 0.340454 | 0.305162 | 0.925193 | 0.144194 | 0.495427 | 0.652678 | 2.863107 |
| TGvd | 0.272115 | 0.230923 | 1.170049 | 0 | 0.70556 | 0.766806 | 3.145453 |
| TGvg | 0.245644 | 0.275094 | 0.839797 | 0.150492 | 0.682917 | 0.603392 | 2.797336 |
| TH | 0.236237 | 0.183019 | 0.597376 | 0 | 0.544954 | 0.605147 | 2.166732 |
| TPO | 0.186205 | 0.224965 | 0.872226 | 0.168276 | 0.3697 | 0.483167 | 2.30454 |
| Tpt | 0.195931 | 0.201281 | 0.668122 | 0.181308 | 0.362726 | 0.387184 | 1.996554 |
| V1 | 0.173575 | 0.139827 | 0.351123 | 0.506209 | 0.160355 | 0.232468 | 1.563556 |
| V2 | 0.161243 | 0.172444 | 0.568535 | 0.148259 | 0.28878 | 0.344185 | 1.683446 |
| V3A | 0.154809 | 0.203265 | 0.625763 | 0.15376 | 0.305252 | 0.328206 | 1.771054 |
| V3d | 0.191741 | 0.19594 | 0.61669 | 0.178648 | 0.288451 | 0.316121 | 1.787591 |
| V3v | 0.180664 | 0.167297 | 0.639161 | 0.150007 | 0.292774 | 0.317127 | 1.747029 |
| V4 | 0.175929 | 0.192766 | 0.665459 | 0.190693 | 0.369131 | 0.588775 | 2.182754 |
| V4t | 0.169372 | 0.207887 | 0.829174 | 0.226331 | 0.476425 | 0.593473 | 2.502661 |
| V4v | 0.217378 | 0.19527 | 0.693806 | 0.180567 | 0.351833 | 0.444395 | 2.083249 |
| V6 | 0.190785 | 0.175651 | 0.531007 | 0.137382 | 0.261788 | 0.330865 | 1.627478 |
| V6Ad | 0.216526 | 0.186371 | 0.778767 | 0.167623 | 0.434214 | 0.50044 | 2.28394 |
| V6Av | 0.183045 | 0.164967 | 0.585577 | 0.110673 | 0.308966 | 0.305882 | 1.659111 |
| VIP | 0.251652 | 0.223797 | 0.638323 | 0.140249 | 0.231837 | 0.273967 | 1.759826 |
| v23b | 0.197133 | 0.199312 | 0.681513 | 0.168154 | 0.443907 | 0.562231 | 2.252251 |

**Table S3. Regional distribution of retrograde-labeled neurons projecting to area 23 based on M132 parcellation**

| **RegionName** | **Hemisphere** | **Neuron Number** | **Layer I** | **Layer II** | **Layer III** | **Layer IV** | **Layer V** | **Layer VI** |
| --- | --- | --- | --- | --- | --- | --- | --- | --- |
| 23 | Ipsilateral | 29326 | 457 | 3671 | 13969 | 2132 | 5648 | 3449 |
| 23 | Contralateral | 8277 | 73 | 810 | 4571 | 616 | 1542 | 665 |
| 29/30 | Ipsilateral | 5752 | 0 | 124 | 1130 | 144 | 3356 | 998 |
| 7A | Ipsilateral | 1459 | 20 | 88 | 582 | 152 | 401 | 216 |
| TF/TH | Ipsilateral | 1403 | 0 | 0 | 562 | 151 | 356 | 334 |
| DP | Ipsilateral | 1342 | 0 | 0 | 355 | 172 | 368 | 447 |
| 29/30 | Contralateral | 1340 | 0 | 134 | 62 | 2 | 783 | 359 |
| 46d | Ipsilateral | 1244 | 0 | 1 | 377 | 212 | 386 | 268 |
| 8B | Ipsilateral | 969 | 2 | 43 | 410 | 0 | 247 | 267 |
| V6A | Ipsilateral | 773 | 0 | 1 | 510 | 62 | 100 | 100 |
| PIP | Ipsilateral | 746 | 0 | 0 | 231 | 94 | 249 | 172 |
| 31 | Ipsilateral | 733 | 3 | 24 | 381 | 77 | 192 | 56 |
| V2fpLF | Ipsilateral | 578 | 0 | 2 | 222 | 87 | 192 | 75 |
| TEpv | Ipsilateral | 430 | 0 | 1 | 42 | 52 | 172 | 163 |
| 46v | Ipsilateral | 411 | 12 | 82 | 275 | 24 | 16 | 2 |
| 7m | Ipsilateral | 397 | 0 | 1 | 231 | 47 | 102 | 16 |
| DP | Contralateral | 356 | 0 | 0 | 65 | 60 | 151 | 80 |
| PIP | Contralateral | 326 | 4 | 3 | 117 | 33 | 105 | 64 |
| 9 | Ipsilateral | 265 | 0 | 0 | 88 | 15 | 61 | 101 |
| PGa | Ipsilateral | 251 | 10 | 8 | 80 | 19 | 24 | 110 |
| Pro.St. | Ipsilateral | 211 | 0 | 3 | 35 | 23 | 127 | 23 |
| LIP | Ipsilateral | 186 | 25 | 20 | 67 | 27 | 43 | 4 |
| ENTO | Ipsilateral | 183 | 0 | 0 | 9 | 0 | 86 | 88 |
| V2fpLF | Contralateral | 148 | 0 | 0 | 79 | 46 | 23 | 0 |
| 8m | Ipsilateral | 127 | 0 | 1 | 45 | 17 | 32 | 32 |
| 24b | Ipsilateral | 106 | 0 | 0 | 18 | 0 | 63 | 25 |
| 9/46d | Ipsilateral | 81 | 0 | 0 | 30 | 14 | 17 | 20 |
| F7 | Ipsilateral | 73 | 0 | 0 | 41 | 0 | 4 | 28 |
| 7A | Contralateral | 72 | 0 | 1 | 40 | 6 | 20 | 5 |
| 11 | Ipsilateral | 69 | 0 | 0 | 10 | 15 | 21 | 23 |
| 31 | Contralateral | 66 | 0 | 1 | 50 | 6 | 5 | 4 |
| 24a | Ipsilateral | 59 | 0 | 0 | 7 | 0 | 32 | 20 |
| 10 | Ipsilateral | 57 | 0 | 0 | 23 | 4 | 14 | 16 |
| LIP | Contralateral | 54 | 0 | 10 | 27 | 10 | 6 | 1 |
| STPi | Ipsilateral | 43 | 0 | 0 | 5 | 1 | 1 | 36 |
| 12 | Ipsilateral | 41 | 0 | 0 | 0 | 5 | 11 | 25 |
| STPr | Ipsilateral | 36 | 0 | 0 | 4 | 3 | 9 | 20 |
| STPc | Ipsilateral | 31 | 0 | 0 | 5 | 2 | 10 | 14 |
| TF/TH | Contralateral | 24 | 0 | 0 | 6 | 5 | 8 | 5 |
| 46v | Contralateral | 20 | 0 | 3 | 13 | 2 | 2 | 0 |
| 8B | Contralateral | 20 | 0 | 0 | 7 | 0 | 7 | 6 |
| V2fpUF | Ipsilateral | 16 | 0 | 1 | 10 | 1 | 3 | 1 |
| POLE | Ipsilateral | 15 | 0 | 0 | 0 | 0 | 1 | 14 |
| Pro.St. | Contralateral | 14 | 0 | 0 | 0 | 6 | 6 | 2 |
| 24b | Contralateral | 14 | 0 | 0 | 2 | 0 | 6 | 6 |
| 14 | Ipsilateral | 13 | 0 | 0 | 1 | 0 | 8 | 4 |
| 8m | Contralateral | 13 | 0 | 0 | 7 | 2 | 2 | 2 |
| PBr | Ipsilateral | 13 | 0 | 0 | 0 | 0 | 7 | 6 |
| 32 | Ipsilateral | 11 | 0 | 0 | 2 | 0 | 5 | 4 |
| STPc | Contralateral | 10 | 0 | 0 | 1 | 2 | 6 | 1 |
| V6A | Contralateral | 10 | 0 | 0 | 1 | 2 | 5 | 2 |
| 13 | Ipsilateral | 9 | 0 | 0 | 2 | 0 | 2 | 5 |
| 46d | Contralateral | 7 | 0 | 0 | 0 | 2 | 4 | 1 |
| ENTO | Contralateral | 7 | 0 | 0 | 1 | 0 | 3 | 3 |
| 7m | Contralateral | 7 | 0 | 0 | 5 | 0 | 0 | 2 |
| 45A | Ipsilateral | 6 | 0 | 0 | 2 | 2 | 1 | 1 |
| 8r | Ipsilateral | 6 | 0 | 0 | 2 | 0 | 1 | 3 |
| PERI | Ipsilateral | 6 | 0 | 0 | 0 | 0 | 1 | 5 |
| V6 | Ipsilateral | 6 | 0 | 0 | 3 | 1 | 2 | 0 |
| V4LF | Contralateral | 5 | 0 | 0 | 0 | 0 | 0 | 5 |
| 9/46d | Contralateral | 5 | 0 | 0 | 2 | 0 | 0 | 3 |
| PGa | Contralateral | 5 | 0 | 0 | 1 | 0 | 0 | 4 |
| STPi | Contralateral | 5 | 0 | 0 | 0 | 0 | 0 | 5 |
| V4LF | Ipsilateral | 4 | 0 | 0 | 0 | 1 | 0 | 3 |
| FST | Ipsilateral | 4 | 0 | 0 | 3 | 1 | 0 | 0 |
| TEpv | Contralateral | 4 | 0 | 0 | 0 | 0 | 0 | 4 |
| 9 | Contralateral | 3 | 0 | 0 | 0 | 0 | 1 | 2 |
| INS | Ipsilateral | 3 | 0 | 0 | 0 | 0 | 0 | 3 |
| MST | Ipsilateral | 3 | 0 | 0 | 0 | 0 | 2 | 1 |
| TPt | Ipsilateral | 3 | 0 | 0 | 0 | 0 | 0 | 3 |
| 24c | Ipsilateral | 3 | 0 | 0 | 0 | 0 | 3 | 0 |
| INS | Contralateral | 2 | 0 | 0 | 0 | 0 | 0 | 2 |
| 12 | Contralateral | 2 | 0 | 0 | 0 | 0 | 0 | 2 |
| SII | Contralateral | 2 | 0 | 0 | 0 | 0 | 1 | 1 |
| PBr | Contralateral | 2 | 0 | 0 | 0 | 0 | 0 | 2 |
| 24a | Contralateral | 2 | 0 | 0 | 0 | 0 | 1 | 1 |
| 32 | Contralateral | 2 | 0 | 0 | 0 | 0 | 1 | 1 |
| 25 | Ipsilateral | 1 | 0 | 0 | 0 | 0 | 1 | 0 |
| 11 | Contralateral | 1 | 0 | 0 | 0 | 0 | 0 | 1 |
| 13 | Contralateral | 1 | 0 | 0 | 0 | 0 | 1 | 0 |
| 14 | Contralateral | 1 | 0 | 0 | 0 | 0 | 1 | 0 |
| 44 | Contralateral | 1 | 0 | 0 | 1 | 0 | 0 | 0 |
| 9/46v | Ipsilateral | 1 | 0 | 0 | 1 | 0 | 0 | 0 |
| F1 | Contralateral | 1 | 0 | 0 | 1 | 0 | 0 | 0 |
| F2 | Ipsilateral | 1 | 0 | 0 | 1 | 0 | 0 | 0 |
| F3 | Ipsilateral | 1 | 0 | 0 | 0 | 0 | 1 | 0 |
| F7 | Contralateral | 1 | 0 | 0 | 0 | 0 | 0 | 1 |
| TEa/ma | Ipsilateral | 1 | 0 | 0 | 0 | 0 | 0 | 1 |
| Pi | Ipsilateral | 1 | 0 | 0 | 0 | 1 | 0 | 0 |
| 7B | Contralateral | 1 | 0 | 0 | 0 | 0 | 0 | 1 |
| MIP | Ipsilateral | 1 | 0 | 0 | 0 | 0 | 0 | 1 |
| MIP | Contralateral | 1 | 0 | 0 | 1 | 0 | 0 | 0 |
| OPRO | Ipsilateral | 1 | 0 | 0 | 0 | 0 | 0 | 1 |
| PBc | Ipsilateral | 1 | 0 | 0 | 0 | 0 | 1 | 0 |
| 24d | Ipsilateral | 1 | 0 | 0 | 0 | 0 | 0 | 1 |
| V1fpLF | Ipsilateral | 1 | 0 | 0 | 1 | 0 | 0 | 0 |

**Table S4. Regional distribution of retrograde-labeled neurons projecting to area 23 based on SARM parcellation**

| **RegionName** | **Hemisphere** | **Neuron Number** |
| --- | --- | --- |
| Cl | Ipsilateral | 1519 |
| MLThal | Ipsilateral | 343 |
| LP | Ipsilateral | 303 |
| f | Ipsilateral | 247 |
| LD | Ipsilateral | 183 |
| DStr | Ipsilateral | 182 |
| ILThal | Ipsilateral | 162 |
| Pul | Ipsilateral | 98 |
| HF | Ipsilateral | 78 |
| A+ | Ipsilateral | 76 |
| SDBR | Ipsilateral | 74 |
| MD | Ipsilateral | 60 |
| MLThal | Contralateral | 40 |
| DTHy | Ipsilateral | 34 |
| lpAmy | Ipsilateral | 24 |
| PNThal | Ipsilateral | 19 |
| DA_Mid | Ipsilateral | 18 |
| HF | Contralateral | 17 |
| ZI-H | Ipsilateral | 15 |
| vpAmy | Ipsilateral | 14 |
| Pd | Ipsilateral | 14 |
| RtMid | Ipsilateral | 14 |
| DPHy | Ipsilateral | 12 |
| raphe_Mid | Contralateral | 12 |
| DTHy | Contralateral | 10 |
| raphe_Mid | Ipsilateral | 10 |
| CbThal | Ipsilateral | 9 |
| BR | Ipsilateral | 7 |
| ZI-H | Contralateral | 7 |
| Co | Ipsilateral | 7 |
| Cl | Contralateral | 5 |
| PAGR | Ipsilateral | 5 |
| R+ | Ipsilateral | 5 |
| mAmy | Ipsilateral | 4 |
| VPHy | Ipsilateral | 4 |
| pdAmy | Ipsilateral | 3 |
| opt | Ipsilateral | 3 |
| SpThal | Ipsilateral | 3 |
| TgMid | Ipsilateral | 3 |
| OC+ | Ipsilateral | 3 |
| DPHy | Contralateral | 2 |
| DA_Mid | Contralateral | 2 |
| lpAmy | Contralateral | 1 |
| DStr | Contralateral | 1 |
| SDBR | Contralateral | 1 |
| opt | Contralateral | 1 |
| VTHy | Ipsilateral | 1 |
| VPHy | Contralateral | 1 |
| ILThal | Contralateral | 1 |
| SpThal | Contralateral | 1 |
| PrT | Ipsilateral | 1 |
| OC+ | Contralateral | 1 |
| R+ | Contralateral | 1 |
| PBC | Ipsilateral | 1 |
| VCbCx | Ipsilateral | 1 |

**Table S5. Regional distribution of retrograde-labeled neurons projecting to area 23 based on D99 parcellation**

| **RegionName** | **Hemisphere** | **Neuron Number** | **Layer I** | **Layer II** | **Layer III** | **Layer IV** | **Layer V** | **Layer VI** |
| --- | --- | --- | --- | --- | --- | --- | --- | --- |
| v23b | Ipsilateral | 18650 | 374 | 3154 | 9694 | 1136 | 2953 | 1339 |
| v23b | Contralateral | 6115 | 61 | 690 | 3911 | 470 | 807 | 176 |
| V2 | Ipsilateral | 4240 | 55 | 216 | 1661 | 474 | 928 | 906 |
| 30 | Ipsilateral | 4205 | 0 | 61 | 1275 | 96 | 2021 | 752 |
| 23a | Ipsilateral | 2553 | 0 | 34 | 1171 | 197 | 673 | 478 |
| 29 | Ipsilateral | 2234 | 0 | 108 | 34 | 0 | 1640 | 452 |
| 7a_(Opt/PG) | Ipsilateral | 2181 | 4 | 54 | 842 | 268 | 609 | 404 |
| 23b | Ipsilateral | 1988 | 3 | 43 | 755 | 127 | 657 | 403 |
| 7m_(PGm) | Ipsilateral | 1901 | 0 | 3 | 1189 | 324 | 315 | 70 |
| 30 | Contralateral | 1285 | 7 | 74 | 410 | 11 | 490 | 293 |
| V2_or_v23b? | Ipsilateral | 1099 | 4 | 64 | 261 | 136 | 372 | 262 |
| 9d | Ipsilateral | 1054 | 1 | 31 | 445 | 17 | 261 | 299 |
| V4 | Ipsilateral | 994 | 0 | 0 | 199 | 126 | 267 | 402 |
| 46d | Ipsilateral | 924 | 0 | 0 | 280 | 174 | 259 | 211 |
| 29 | Contralateral | 834 | 0 | 112 | 4 | 0 | 527 | 191 |
| V3v | Ipsilateral | 773 | 0 | 0 | 468 | 101 | 171 | 33 |
| 46f | Ipsilateral | 677 | 9 | 62 | 307 | 74 | 158 | 67 |
| V2 | Contralateral | 613 | 5 | 16 | 190 | 107 | 220 | 75 |
| TEpv | Ipsilateral | 469 | 0 | 0 | 45 | 64 | 184 | 176 |
| V2? | Ipsilateral | 379 | 0 | 65 | 146 | 42 | 80 | 46 |
| V4 | Contralateral | 366 | 0 | 0 | 55 | 44 | 157 | 110 |
| V2_or_v23b? | Contralateral | 362 | 1 | 0 | 39 | 42 | 158 | 122 |
| LIPd | Ipsilateral | 357 | 33 | 53 | 117 | 34 | 102 | 18 |
| TF | Ipsilateral | 307 | 0 | 0 | 11 | 9 | 88 | 199 |
| 7a_(Opt/PG) | Contralateral | 256 | 0 | 1 | 87 | 41 | 87 | 40 |
| 31 | Ipsilateral | 245 | 0 | 11 | 152 | 33 | 39 | 10 |
| TFO | Ipsilateral | 242 | 0 | 0 | 74 | 29 | 79 | 60 |
| 8Bs | Ipsilateral | 202 | 1 | 13 | 88 | 8 | 51 | 41 |
| LOP | Ipsilateral | 197 | 0 | 0 | 76 | 18 | 83 | 20 |
| 23b | Contralateral | 179 | 0 | 0 | 51 | 9 | 55 | 64 |
| PGa | Ipsilateral | 160 | 10 | 9 | 55 | 11 | 12 | 63 |
| 23a | Contralateral | 158 | 0 | 1 | 50 | 8 | 33 | 66 |
| TPO | Ipsilateral | 158 | 0 | 0 | 33 | 10 | 25 | 90 |
| 24b' | Ipsilateral | 137 | 0 | 0 | 39 | 2 | 42 | 54 |
| V2? | Contralateral | 133 | 0 | 0 | 38 | 21 | 47 | 27 |
| 46v | Ipsilateral | 129 | 3 | 21 | 98 | 2 | 4 | 1 |
| LOP | Contralateral | 126 | 4 | 3 | 73 | 14 | 29 | 3 |
| EC | Ipsilateral | 124 | 0 | 0 | 9 | 0 | 63 | 52 |
| 8Bd | Ipsilateral | 109 | 0 | 0 | 32 | 0 | 10 | 67 |
| 24b | Ipsilateral | 99 | 0 | 0 | 12 | 0 | 69 | 18 |
| 11l | Ipsilateral | 81 | 0 | 0 | 9 | 16 | 21 | 35 |
| 8Ad | Ipsilateral | 78 | 0 | 0 | 32 | 9 | 24 | 13 |
| v23b? | Ipsilateral | 64 | 0 | 28 | 36 | 0 | 0 | 0 |
| LIPd | Contralateral | 64 | 0 | 10 | 34 | 9 | 9 | 2 |
| ECL | Ipsilateral | 51 | 0 | 0 | 1 | 0 | 16 | 34 |
| 31 | Contralateral | 49 | 0 | 0 | 44 | 2 | 3 | 0 |
| TAa | Ipsilateral | 42 | 0 | 0 | 5 | 3 | 10 | 24 |
| V4v | Ipsilateral | 28 | 0 | 1 | 4 | 1 | 8 | 14 |
| 7m_(PGm) | Contralateral | 28 | 0 | 0 | 16 | 5 | 2 | 5 |
| 9d | Contralateral | 18 | 0 | 0 | 3 | 0 | 7 | 8 |
| 24a' | Ipsilateral | 17 | 0 | 0 | 7 | 0 | 9 | 1 |
| TPO | Contralateral | 15 | 0 | 0 | 1 | 2 | 3 | 9 |
| 24a | Ipsilateral | 15 | 0 | 0 | 2 | 0 | 8 | 5 |
| V3v | Contralateral | 14 | 0 | 0 | 4 | 5 | 5 | 0 |
| 11m | Ipsilateral | 14 | 0 | 0 | 1 | 3 | 8 | 2 |
| 9m | Ipsilateral | 13 | 0 | 0 | 2 | 1 | 4 | 6 |
| EI | Ipsilateral | 13 | 0 | 0 | 0 | 0 | 11 | 2 |
| 46v | Contralateral | 13 | 0 | 0 | 10 | 2 | 1 | 0 |
| 23c | Ipsilateral | 13 | 0 | 1 | 8 | 0 | 1 | 3 |
| 8Ad | Contralateral | 13 | 0 | 0 | 6 | 2 | 2 | 3 |
| 10mr | Ipsilateral | 11 | 0 | 0 | 4 | 0 | 2 | 5 |
| IPa | Ipsilateral | 11 | 0 | 0 | 4 | 1 | 0 | 6 |
| TH | Ipsilateral | 10 | 0 | 0 | 2 | 0 | 4 | 4 |
| 12m | Ipsilateral | 10 | 0 | 0 | 0 | 1 | 2 | 7 |
| 46f | Contralateral | 9 | 0 | 3 | 3 | 1 | 1 | 1 |
| STGr | Ipsilateral | 9 | 0 | 0 | 0 | 0 | 2 | 7 |
| 24b' | Contralateral | 9 | 0 | 0 | 2 | 0 | 2 | 5 |
| Tpt | Ipsilateral | 8 | 0 | 0 | 0 | 1 | 3 | 4 |
| 13l | Ipsilateral | 8 | 0 | 0 | 2 | 0 | 2 | 4 |
| 8Bs | Contralateral | 8 | 0 | 0 | 7 | 0 | 1 | 0 |
| 13m | Ipsilateral | 7 | 0 | 0 | 0 | 0 | 7 | 0 |
| 46d | Contralateral | 7 | 0 | 0 | 0 | 1 | 4 | 2 |
| 12r | Ipsilateral | 6 | 0 | 0 | 0 | 1 | 4 | 1 |
| V1 | Ipsilateral | 5 | 0 | 1 | 3 | 0 | 1 | 0 |
| 24b | Contralateral | 5 | 0 | 0 | 1 | 0 | 4 | 0 |
| ECL | Contralateral | 5 | 0 | 0 | 1 | 0 | 3 | 1 |
| TEpv | Contralateral | 5 | 0 | 0 | 0 | 0 | 0 | 5 |
| 45a | Ipsilateral | 5 | 0 | 0 | 2 | 2 | 0 | 1 |
| ELc | Ipsilateral | 5 | 0 | 0 | 0 | 0 | 1 | 4 |
| 35 | Ipsilateral | 4 | 0 | 0 | 0 | 0 | 0 | 4 |
| 24c | Ipsilateral | 4 | 0 | 0 | 0 | 0 | 3 | 1 |
| TF | Contralateral | 4 | 0 | 0 | 0 | 0 | 0 | 4 |
| TGsts | Ipsilateral | 3 | 0 | 0 | 0 | 0 | 0 | 3 |
| Id | Ipsilateral | 3 | 0 | 0 | 0 | 0 | 0 | 3 |
| 8Bm | Ipsilateral | 3 | 0 | 0 | 0 | 0 | 0 | 3 |
| 36c | Ipsilateral | 3 | 0 | 0 | 0 | 0 | 1 | 2 |
| 32 | Ipsilateral | 3 | 0 | 0 | 1 | 0 | 1 | 1 |
| TFO | Contralateral | 3 | 0 | 0 | 1 | 0 | 2 | 0 |
| TGdd | Ipsilateral | 3 | 0 | 0 | 0 | 0 | 0 | 3 |
| EC | Contralateral | 3 | 0 | 0 | 0 | 0 | 1 | 2 |
| MIP | Ipsilateral | 3 | 0 | 0 | 0 | 0 | 2 | 1 |
| F7 | Ipsilateral | 3 | 0 | 0 | 0 | 0 | 0 | 3 |
| TGa | Ipsilateral | 3 | 0 | 0 | 0 | 0 | 0 | 3 |
| F2_(6DR/6DC) | Ipsilateral | 3 | 0 | 0 | 2 | 0 | 1 | 0 |
| 24c' | Ipsilateral | 3 | 0 | 0 | 0 | 0 | 1 | 2 |
| Id | Contralateral | 2 | 0 | 0 | 0 | 0 | 0 | 2 |
| Iai | Ipsilateral | 2 | 0 | 0 | 0 | 0 | 0 | 2 |
| 13b | Ipsilateral | 2 | 0 | 0 | 0 | 0 | 0 | 2 |
| TEav | Ipsilateral | 2 | 0 | 0 | 0 | 0 | 0 | 2 |
| V6Av | Ipsilateral | 2 | 0 | 0 | 1 | 0 | 1 | 0 |
| SII | Contralateral | 2 | 0 | 0 | 0 | 0 | 1 | 1 |
| RPB | Ipsilateral | 2 | 0 | 0 | 0 | 0 | 0 | 2 |
| v23b? | Contralateral | 2 | 0 | 0 | 2 | 0 | 0 | 0 |
| Tpt | Contralateral | 2 | 0 | 0 | 0 | 0 | 2 | 0 |
| MST | Ipsilateral | 2 | 0 | 0 | 0 | 0 | 1 | 1 |
| STGr | Contralateral | 2 | 0 | 0 | 0 | 0 | 0 | 2 |
| F6 | Ipsilateral | 2 | 0 | 0 | 0 | 0 | 0 | 2 |
| PrCO | Ipsilateral | 2 | 0 | 0 | 0 | 0 | 0 | 2 |
| V6Ad | Ipsilateral | 2 | 0 | 0 | 1 | 0 | 1 | 0 |
| 12o | Ipsilateral | 2 | 0 | 0 | 0 | 0 | 0 | 2 |
| 24a | Contralateral | 2 | 0 | 0 | 0 | 0 | 1 | 1 |
| TGvd | Ipsilateral | 1 | 0 | 0 | 0 | 0 | 0 | 1 |
| 24c' | Contralateral | 1 | 0 | 0 | 0 | 0 | 0 | 1 |
| 13b | Contralateral | 1 | 0 | 0 | 0 | 0 | 1 | 0 |
| CPB | Ipsilateral | 1 | 0 | 0 | 0 | 0 | 1 | 0 |
| 13l | Contralateral | 1 | 0 | 0 | 0 | 0 | 1 | 0 |
| VIP | Ipsilateral | 1 | 0 | 0 | 0 | 0 | 1 | 0 |
| LIPv | Contralateral | 1 | 0 | 0 | 0 | 1 | 0 | 0 |
| 12 | Contralateral | 1 | 0 | 0 | 0 | 0 | 0 | 1 |
| 12m | Contralateral | 1 | 0 | 0 | 0 | 0 | 0 | 1 |
| F1_(4) | Contralateral | 1 | 0 | 0 | 1 | 0 | 0 | 0 |
| 13m | Contralateral | 1 | 0 | 0 | 0 | 0 | 0 | 1 |
| F3 | Ipsilateral | 1 | 0 | 0 | 0 | 0 | 1 | 0 |
| Ial | Ipsilateral | 1 | 0 | 0 | 0 | 0 | 0 | 1 |
| 23c | Contralateral | 1 | 0 | 0 | 1 | 0 | 0 | 0 |
| 44 | Contralateral | 1 | 0 | 0 | 1 | 0 | 0 | 0 |
| TEm | Ipsilateral | 1 | 0 | 0 | 0 | 0 | 0 | 1 |
| 24c | Contralateral | 1 | 0 | 0 | 0 | 0 | 1 | 0 |
| MST | Contralateral | 1 | 0 | 0 | 0 | 0 | 1 | 0 |
| RM | Ipsilateral | 1 | 0 | 0 | 0 | 1 | 0 | 0 |
| TGdg | Ipsilateral | 1 | 0 | 0 | 0 | 0 | 0 | 1 |
| F5_(6Va/6Vb) | Ipsilateral | 1 | 0 | 0 | 0 | 0 | 1 | 0 |
| 5_(PEa) | Contralateral | 1 | 0 | 0 | 1 | 0 | 0 | 0 |
| PGa | Contralateral | 1 | 0 | 0 | 1 | 0 | 0 | 0 |
| 12o | Contralateral | 1 | 0 | 0 | 0 | 0 | 0 | 1 |
| TAa | Contralateral | 1 | 0 | 0 | 0 | 0 | 0 | 1 |
| F2_(6DR/6DC) | Contralateral | 1 | 0 | 0 | 0 | 0 | 0 | 1 |
| TGvg | Ipsilateral | 1 | 0 | 0 | 0 | 0 | 0 | 1 |
| ELr | Ipsilateral | 1 | 0 | 0 | 0 | 0 | 0 | 1 |

**Table S6. Regional distribution of retrograde-labeled neurons from claustrum based on M132 parcellation**

| **RegionName** | **Hemisphere** | **Neuron Number** | **Layer I** | **Layer II** | **Layer III** | **Layer IV** | **Layer V** | **Layer VI** |
| --- | --- | --- | --- | --- | --- | --- | --- | --- |
| Pi | Ipsilateral | 2290 | 10 | 63 | 1307 | 0 | 855 | 55 |
| LIP | Ipsilateral | 2110 | 4 | 25 | 156 | 216 | 1054 | 655 |
| 23 | Ipsilateral | 1911 | 1 | 7 | 203 | 63 | 543 | 1094 |
| PIP | Ipsilateral | 1438 | 27 | 31 | 293 | 94 | 483 | 510 |
| MTp | Ipsilateral | 1370 | 7 | 17 | 257 | 119 | 467 | 503 |
| PGa | Ipsilateral | 1179 | 9 | 21 | 383 | 115 | 287 | 364 |
| V6A | Ipsilateral | 990 | 4 | 6 | 67 | 59 | 344 | 510 |
| INS | Ipsilateral | 875 | 4 | 24 | 286 | 80 | 207 | 274 |
| TF/TH | Ipsilateral | 839 | 0 | 0 | 23 | 28 | 460 | 328 |
| STPi | Ipsilateral | 800 | 2 | 14 | 179 | 70 | 175 | 360 |
| V4LF | Ipsilateral | 662 | 30 | 43 | 185 | 75 | 118 | 211 |
| DP | Ipsilateral | 642 | 12 | 43 | 161 | 32 | 127 | 267 |
| V4t | Ipsilateral | 552 | 21 | 26 | 199 | 95 | 166 | 45 |
| 7A | Ipsilateral | 521 | 2 | 1 | 72 | 40 | 148 | 258 |
| 8l | Ipsilateral | 519 | 0 | 7 | 331 | 21 | 47 | 113 |
| 8m | Ipsilateral | 518 | 4 | 13 | 225 | 56 | 88 | 132 |
| VIP | Ipsilateral | 512 | 2 | 7 | 52 | 59 | 208 | 184 |
| TEpv | Ipsilateral | 505 | 1 | 9 | 87 | 31 | 102 | 275 |
| TEOm | Ipsilateral | 382 | 2 | 10 | 55 | 31 | 164 | 120 |
| 31 | Ipsilateral | 376 | 0 | 1 | 35 | 26 | 161 | 153 |
| MB | Ipsilateral | 361 | 17 | 29 | 170 | 31 | 102 | 12 |
| FST | Ipsilateral | 351 | 2 | 5 | 69 | 30 | 111 | 134 |
| MST | Ipsilateral | 350 | 1 | 4 | 34 | 52 | 149 | 110 |
| STPr | Ipsilateral | 335 | 12 | 23 | 109 | 26 | 54 | 111 |
| 7m | Ipsilateral | 324 | 0 | 0 | 22 | 2 | 53 | 247 |
| TEO | Ipsilateral | 298 | 3 | 9 | 63 | 31 | 79 | 113 |
| 8B | Ipsilateral | 287 | 3 | 9 | 152 | 0 | 32 | 91 |
| STPc | Ipsilateral | 281 | 2 | 3 | 90 | 34 | 40 | 112 |
| V4c | Ipsilateral | 281 | 2 | 2 | 53 | 32 | 94 | 98 |
| MTc | Ipsilateral | 270 | 4 | 5 | 80 | 60 | 107 | 14 |
| V6 | Ipsilateral | 267 | 0 | 0 | 13 | 41 | 122 | 91 |
| IPa | Ipsilateral | 262 | 0 | 7 | 139 | 25 | 63 | 28 |
| ENTO | Ipsilateral | 205 | 0 | 0 | 2 | 0 | 26 | 177 |
| TEa/mp | Ipsilateral | 202 | 3 | 9 | 66 | 27 | 69 | 28 |
| V3A | Ipsilateral | 185 | 1 | 8 | 101 | 19 | 50 | 6 |
| TEpd | Ipsilateral | 181 | 0 | 0 | 3 | 8 | 14 | 156 |
| 45B | Ipsilateral | 171 | 3 | 3 | 102 | 4 | 22 | 37 |
| 12 | Ipsilateral | 167 | 0 | 1 | 26 | 10 | 40 | 90 |
| POLE | Ipsilateral | 154 | 1 | 6 | 29 | 3 | 44 | 71 |
| 24b | Ipsilateral | 146 | 0 | 0 | 23 | 0 | 60 | 63 |
| 29/30 | Ipsilateral | 145 | 0 | 3 | 55 | 3 | 31 | 53 |
| V4UF | Ipsilateral | 136 | 0 | 1 | 13 | 8 | 38 | 76 |
| PBc | Ipsilateral | 135 | 0 | 0 | 3 | 18 | 53 | 61 |
| 45A | Ipsilateral | 131 | 0 | 0 | 17 | 11 | 30 | 73 |
| PBr | Ipsilateral | 126 | 1 | 2 | 13 | 4 | 31 | 75 |
| PERI | Ipsilateral | 120 | 0 | 2 | 6 | 0 | 12 | 100 |
| TEa/ma | Ipsilateral | 110 | 1 | 4 | 36 | 20 | 25 | 24 |
| F4 | Ipsilateral | 106 | 5 | 11 | 70 | 0 | 11 | 9 |
| 46v | Ipsilateral | 104 | 0 | 4 | 48 | 19 | 23 | 10 |
| MB | Contralateral | 102 | 8 | 13 | 80 | 1 | 0 | 0 |
| 8r | Ipsilateral | 101 | 2 | 1 | 12 | 12 | 21 | 53 |
| F2 | Ipsilateral | 99 | 1 | 6 | 39 | 0 | 28 | 25 |
| SII | Ipsilateral | 91 | 0 | 2 | 37 | 5 | 34 | 13 |
| 46d | Ipsilateral | 88 | 0 | 0 | 24 | 6 | 12 | 46 |
| Core | Ipsilateral | 85 | 2 | 7 | 41 | 4 | 18 | 13 |
| F7 | Ipsilateral | 84 | 0 | 0 | 2 | 0 | 24 | 58 |
| Pro.St. | Ipsilateral | 77 | 0 | 0 | 20 | 16 | 36 | 5 |
| MIP | Ipsilateral | 71 | 0 | 0 | 11 | 2 | 8 | 50 |
| V2fpLF | Ipsilateral | 69 | 0 | 0 | 11 | 16 | 19 | 23 |
| V2c | Ipsilateral | 69 | 1 | 2 | 34 | 11 | 12 | 9 |
| 13 | Ipsilateral | 68 | 0 | 0 | 5 | 0 | 2 | 61 |
| 9/46v | Ipsilateral | 68 | 0 | 0 | 21 | 6 | 14 | 27 |
| 23 | Contralateral | 68 | 0 | 0 | 16 | 2 | 11 | 39 |
| 9 | Ipsilateral | 67 | 0 | 0 | 9 | 4 | 14 | 40 |
| TPt | Ipsilateral | 63 | 4 | 1 | 10 | 6 | 6 | 36 |
| LB | Ipsilateral | 63 | 0 | 0 | 9 | 11 | 19 | 24 |
| TEav | Ipsilateral | 59 | 0 | 0 | 8 | 1 | 8 | 42 |
| INS | Contralateral | 57 | 0 | 0 | 38 | 11 | 8 | 0 |
| V3pcLF | Ipsilateral | 55 | 0 | 3 | 15 | 8 | 10 | 19 |
| 7A | Contralateral | 53 | 0 | 3 | 14 | 6 | 17 | 13 |
| ENTO | Contralateral | 52 | 0 | 0 | 0 | 0 | 0 | 52 |
| 24c | Ipsilateral | 52 | 0 | 0 | 10 | 0 | 20 | 22 |
| 44 | Ipsilateral | 47 | 2 | 1 | 30 | 0 | 9 | 5 |
| V3fpLF | Ipsilateral | 44 | 0 | 3 | 8 | 3 | 14 | 16 |
| Core | Contralateral | 44 | 1 | 4 | 27 | 6 | 5 | 1 |
| V3c | Ipsilateral | 43 | 0 | 0 | 1 | 0 | 11 | 31 |
| 11 | Ipsilateral | 42 | 0 | 0 | 2 | 1 | 4 | 35 |
| V2pcLF | Ipsilateral | 42 | 1 | 2 | 10 | 11 | 3 | 15 |
| STPi | Contralateral | 39 | 0 | 0 | 16 | 6 | 13 | 4 |
| PGa | Contralateral | 37 | 1 | 1 | 14 | 9 | 8 | 4 |
| 7op | Ipsilateral | 37 | 1 | 2 | 11 | 1 | 5 | 17 |
| LIP | Contralateral | 35 | 0 | 5 | 9 | 1 | 10 | 10 |
| 24a | Ipsilateral | 35 | 0 | 1 | 16 | 0 | 16 | 2 |
| 9/46d | Ipsilateral | 32 | 0 | 1 | 4 | 2 | 4 | 21 |
| 24d | Ipsilateral | 32 | 0 | 0 | 16 | 0 | 7 | 9 |
| V3pcUF | Ipsilateral | 32 | 0 | 0 | 4 | 5 | 8 | 15 |
| F6 | Ipsilateral | 31 | 0 | 0 | 8 | 0 | 5 | 18 |
| 14 | Ipsilateral | 30 | 2 | 5 | 10 | 0 | 5 | 8 |
| AIP | Ipsilateral | 26 | 0 | 0 | 2 | 1 | 19 | 4 |
| V2fpUF | Ipsilateral | 22 | 1 | 1 | 13 | 1 | 1 | 5 |
| Pi | Contralateral | 20 | 0 | 8 | 12 | 0 | 0 | 0 |
| F5 | Ipsilateral | 19 | 0 | 0 | 5 | 0 | 2 | 12 |
| V3fpUF | Ipsilateral | 19 | 4 | 2 | 7 | 0 | 2 | 4 |
| STPc | Contralateral | 18 | 0 | 1 | 10 | 2 | 2 | 3 |
| 24a | Contralateral | 18 | 0 | 0 | 4 | 0 | 11 | 3 |
| 7B | Ipsilateral | 17 | 0 | 0 | 4 | 2 | 4 | 7 |
| Gu | Ipsilateral | 17 | 0 | 3 | 4 | 0 | 8 | 2 |
| 24b | Contralateral | 17 | 0 | 0 | 2 | 0 | 4 | 11 |
| TEad | Ipsilateral | 15 | 0 | 0 | 1 | 1 | 1 | 12 |
| MST | Contralateral | 15 | 0 | 0 | 9 | 1 | 4 | 1 |
| 5 | Ipsilateral | 15 | 0 | 0 | 1 | 0 | 3 | 11 |
| 29/30 | Contralateral | 15 | 0 | 0 | 7 | 0 | 5 | 3 |
| V2pcUF | Ipsilateral | 14 | 0 | 0 | 3 | 5 | 4 | 2 |
| PBr | Contralateral | 14 | 0 | 0 | 0 | 2 | 12 | 0 |
| OPRO | Ipsilateral | 13 | 0 | 1 | 3 | 0 | 0 | 9 |
| LB | Contralateral | 13 | 0 | 0 | 6 | 3 | 4 | 0 |
| PBc | Contralateral | 12 | 0 | 0 | 2 | 1 | 9 | 0 |
| 32 | Ipsilateral | 11 | 0 | 0 | 2 | 0 | 6 | 3 |
| TPt | Contralateral | 10 | 0 | 0 | 2 | 1 | 4 | 3 |
| 10 | Ipsilateral | 10 | 0 | 0 | 1 | 0 | 1 | 8 |
| 8B | Contralateral | 10 | 0 | 0 | 4 | 0 | 2 | 4 |
| PERI | Contralateral | 9 | 0 | 0 | 0 | 0 | 0 | 9 |
| STPr | Contralateral | 9 | 1 | 1 | 5 | 0 | 0 | 2 |
| POLE | Contralateral | 8 | 0 | 0 | 1 | 1 | 5 | 1 |
| 24c | Contralateral | 7 | 0 | 0 | 1 | 0 | 4 | 2 |
| 31 | Contralateral | 7 | 0 | 0 | 1 | 0 | 4 | 2 |
| IPa | Contralateral | 6 | 0 | 0 | 3 | 1 | 2 | 0 |
| DP | Contralateral | 6 | 1 | 2 | 1 | 0 | 2 | 0 |
| F3 | Ipsilateral | 5 | 0 | 0 | 0 | 0 | 2 | 3 |
| 2 | Ipsilateral | 5 | 0 | 0 | 0 | 1 | 1 | 3 |
| 24d | Contralateral | 5 | 0 | 0 | 2 | 0 | 0 | 3 |
| 8m | Contralateral | 4 | 0 | 0 | 0 | 1 | 2 | 1 |
| F7 | Contralateral | 4 | 0 | 0 | 1 | 0 | 1 | 2 |
| TEa/mp | Contralateral | 4 | 0 | 0 | 1 | 0 | 3 | 0 |
| TF/TH | Contralateral | 4 | 0 | 0 | 0 | 0 | 3 | 1 |
| ProM | Ipsilateral | 3 | 0 | 0 | 0 | 0 | 1 | 2 |
| SII | Contralateral | 3 | 0 | 0 | 1 | 1 | 1 | 0 |
| FST | Contralateral | 3 | 0 | 0 | 2 | 0 | 0 | 1 |
| TEpv | Contralateral | 3 | 0 | 0 | 1 | 0 | 1 | 1 |
| 46d | Contralateral | 2 | 0 | 0 | 1 | 0 | 1 | 0 |
| PIP | Contralateral | 2 | 0 | 0 | 0 | 0 | 1 | 1 |
| 32 | Contralateral | 2 | 0 | 0 | 0 | 0 | 2 | 0 |
| 45A | Contralateral | 1 | 0 | 0 | 1 | 0 | 0 | 0 |
| 9 | Contralateral | 1 | 0 | 0 | 0 | 0 | 0 | 1 |
| 9/46v | Contralateral | 1 | 0 | 0 | 1 | 0 | 0 | 0 |
| ProM | Contralateral | 1 | 0 | 0 | 1 | 0 | 0 | 0 |
| F2 | Contralateral | 1 | 0 | 0 | 0 | 0 | 0 | 1 |
| F6 | Contralateral | 1 | 0 | 0 | 0 | 0 | 0 | 1 |
| TEa/ma | Contralateral | 1 | 0 | 0 | 0 | 1 | 0 | 0 |
| 5 | Contralateral | 1 | 0 | 0 | 0 | 0 | 1 | 0 |
| 7B | Contralateral | 1 | 0 | 0 | 0 | 1 | 0 | 0 |
| 7op | Contralateral | 1 | 0 | 0 | 1 | 0 | 0 | 0 |
| VIP | Contralateral | 1 | 0 | 0 | 0 | 0 | 0 | 1 |

**Table S7 Regional distribution of retrograde-labeled neurons from claustrum based on SARM parcellation**

| **RegionName** | **Hemisphere** | **Neuron Number** |
| --- | --- | --- |
| Cl | Ipsilateral | 6419 |
| DStr | Ipsilateral | 973 |
| lpAmy | Ipsilateral | 476 |
| MG | Ipsilateral | 387 |
| Pul | Ipsilateral | 372 |
| vpAmy | Ipsilateral | 253 |
| ILThal | Ipsilateral | 214 |
| HF | Ipsilateral | 90 |
| PrT | Ipsilateral | 79 |
| DLG | Ipsilateral | 75 |
| En | Ipsilateral | 37 |
| MLThal | Ipsilateral | 37 |
| DTHy | Ipsilateral | 30 |
| Co | Ipsilateral | 30 |
| lpAmy | Contralateral | 25 |
| Pd | Ipsilateral | 21 |
| MD | Ipsilateral | 19 |
| SpThal | Ipsilateral | 19 |
| RtMid | Ipsilateral | 12 |
| BR | Ipsilateral | 11 |
| DA_Mid | Ipsilateral | 10 |
| PNThal | Ipsilateral | 9 |
| vpAmy | Contralateral | 7 |
| mAmy | Ipsilateral | 6 |
| LP | Ipsilateral | 5 |
| VPHy | Ipsilateral | 4 |
| HF | Contralateral | 3 |
| stAmy | Ipsilateral | 3 |
| DPHy | Ipsilateral | 3 |
| raphe_Mid | Ipsilateral | 3 |
| DTHy | Contralateral | 2 |
| ZI-H | Ipsilateral | 2 |
| Rt | Ipsilateral | 2 |
| A+ | Ipsilateral | 2 |
| Pul | Contralateral | 2 |
| Hb | Ipsilateral | 2 |
| TgMid | Ipsilateral | 2 |
| Pir | Ipsilateral | 1 |
| DStr | Contralateral | 1 |
| DPHy | Contralateral | 1 |
| LD | Ipsilateral | 1 |
| PAGR | Ipsilateral | 1 |
| raphe_Mid | Contralateral | 1 |
| PBC | Ipsilateral | 1 |

**Table S8. Regional distribution of retrograde-labeled neurons from clsustrum based on D99 parcellation**

| **RegionName** | **Hemisphere** | **Neuron** **Number** | **Layer I** | **Layer II** | **Layer III** | **Layer IV** | **Layer V** | **Layer VI** |
| --- | --- | --- | --- | --- | --- | --- | --- | --- |
| Pi | Ipsilateral | 2147 | 9 | 56 | 1208 | 0 | 816 | 58 |
| V4 | Ipsilateral | 1951 | 30 | 62 | 449 | 201 | 483 | 726 |
| LIPv | Ipsilateral | 1342 | 0 | 12 | 49 | 89 | 743 | 449 |
| TPO | Ipsilateral | 1139 | 11 | 38 | 346 | 108 | 182 | 454 |
| VIP | Ipsilateral | 1097 | 7 | 19 | 181 | 97 | 392 | 401 |
| V2 | Ipsilateral | 874 | 4 | 7 | 97 | 85 | 254 | 427 |
| LIPd | Ipsilateral | 859 | 6 | 8 | 109 | 123 | 394 | 219 |
| MT | Ipsilateral | 844 | 23 | 32 | 235 | 83 | 270 | 201 |
| IPa | Ipsilateral | 815 | 0 | 3 | 301 | 79 | 198 | 234 |
| v23b | Ipsilateral | 763 | 0 | 2 | 47 | 33 | 339 | 342 |
| 7a_(Opt/PG) | Ipsilateral | 714 | 8 | 25 | 110 | 46 | 175 | 350 |
| PGa | Ipsilateral | 707 | 7 | 15 | 207 | 60 | 208 | 210 |
| 23b | Ipsilateral | 702 | 0 | 2 | 89 | 15 | 164 | 432 |
| TEO | Ipsilateral | 681 | 4 | 13 | 121 | 59 | 260 | 224 |
| V3A | Ipsilateral | 657 | 18 | 46 | 261 | 74 | 159 | 99 |
| 8Bs | Ipsilateral | 612 | 0 | 6 | 382 | 41 | 77 | 106 |
| 7m_(PGm) | Ipsilateral | 597 | 0 | 2 | 29 | 40 | 245 | 281 |
| Id | Ipsilateral | 536 | 0 | 10 | 166 | 30 | 158 | 172 |
| MST | Ipsilateral | 528 | 1 | 4 | 62 | 73 | 195 | 193 |
| FST | Ipsilateral | 466 | 6 | 15 | 117 | 82 | 153 | 93 |
| TEpv | Ipsilateral | 418 | 1 | 8 | 68 | 25 | 81 | 235 |
| TAa | Ipsilateral | 398 | 8 | 10 | 80 | 35 | 91 | 174 |
| TFO | Ipsilateral | 379 | 0 | 0 | 10 | 6 | 259 | 104 |
| MIP | Ipsilateral | 354 | 1 | 0 | 39 | 25 | 74 | 215 |
| 45b | Ipsilateral | 337 | 4 | 7 | 208 | 9 | 26 | 83 |
| V4v | Ipsilateral | 336 | 1 | 3 | 44 | 30 | 137 | 121 |
| PIP | Ipsilateral | 324 | 5 | 8 | 61 | 18 | 109 | 123 |
| F2_(6DR/6DC) | Ipsilateral | 288 | 10 | 24 | 154 | 0 | 53 | 47 |
| Ig | Ipsilateral | 281 | 0 | 11 | 124 | 30 | 52 | 64 |
| TF | Ipsilateral | 259 | 0 | 0 | 8 | 2 | 107 | 142 |
| 31 | Ipsilateral | 255 | 0 | 1 | 33 | 14 | 67 | 140 |
| TEpd | Ipsilateral | 221 | 0 | 0 | 5 | 7 | 16 | 193 |
| RTM | Ipsilateral | 216 | 0 | 1 | 121 | 3 | 85 | 6 |
| 8Ad | Ipsilateral | 212 | 0 | 0 | 91 | 23 | 30 | 68 |
| 46d | Ipsilateral | 184 | 0 | 0 | 33 | 11 | 34 | 106 |
| 23a | Ipsilateral | 180 | 0 | 2 | 49 | 4 | 36 | 89 |
| TEa | Ipsilateral | 174 | 2 | 15 | 73 | 34 | 38 | 12 |
| 45a | Ipsilateral | 173 | 0 | 0 | 21 | 17 | 39 | 96 |
| TEav | Ipsilateral | 172 | 0 | 1 | 23 | 7 | 32 | 109 |
| LOP | Ipsilateral | 169 | 3 | 5 | 39 | 11 | 38 | 73 |
| 46v | Ipsilateral | 167 | 2 | 5 | 57 | 25 | 38 | 40 |
| V3v | Ipsilateral | 157 | 4 | 2 | 15 | 22 | 39 | 75 |
| CPB | Ipsilateral | 154 | 0 | 0 | 6 | 16 | 54 | 78 |
| 23c | Ipsilateral | 153 | 1 | 0 | 36 | 5 | 17 | 94 |
| TEm | Ipsilateral | 152 | 2 | 3 | 40 | 13 | 54 | 40 |
| EC | Ipsilateral | 146 | 0 | 0 | 3 | 0 | 8 | 135 |
| CM | Ipsilateral | 140 | 18 | 30 | 65 | 9 | 11 | 7 |
| SII | Ipsilateral | 113 | 0 | 1 | 41 | 11 | 37 | 23 |
| RM | Ipsilateral | 113 | 1 | 4 | 64 | 22 | 21 | 1 |
| 24b' | Ipsilateral | 108 | 0 | 0 | 25 | 0 | 34 | 49 |
| 8Av | Ipsilateral | 105 | 0 | 0 | 30 | 6 | 20 | 49 |
| V3d | Ipsilateral | 103 | 0 | 0 | 20 | 12 | 17 | 54 |
| 30 | Ipsilateral | 102 | 0 | 2 | 32 | 2 | 18 | 48 |
| V2_or_v23b? | Ipsilateral | 91 | 0 | 0 | 6 | 7 | 43 | 35 |
| V6Av | Ipsilateral | 91 | 0 | 1 | 2 | 7 | 51 | 30 |
| STGr | Ipsilateral | 90 | 0 | 1 | 2 | 5 | 23 | 59 |
| F5_(6Va/6Vb) | Ipsilateral | 86 | 4 | 8 | 38 | 0 | 12 | 24 |
| 24b | Ipsilateral | 86 | 0 | 0 | 13 | 0 | 42 | 31 |
| 9d | Ipsilateral | 71 | 0 | 0 | 11 | 0 | 9 | 51 |
| 35 | Ipsilateral | 71 | 0 | 1 | 2 | 0 | 8 | 60 |
| RPB | Ipsilateral | 70 | 0 | 0 | 3 | 2 | 20 | 45 |
| 7a_(Opt/PG) | Contralateral | 70 | 1 | 2 | 12 | 7 | 31 | 17 |
| 13m | Ipsilateral | 69 | 2 | 4 | 16 | 0 | 8 | 39 |
| Tpt | Ipsilateral | 67 | 0 | 0 | 5 | 3 | 13 | 46 |
| F7 | Ipsilateral | 62 | 0 | 0 | 2 | 0 | 18 | 42 |
| RM | Contralateral | 61 | 1 | 10 | 50 | 0 | 0 | 0 |
| V4t | Ipsilateral | 60 | 0 | 0 | 4 | 11 | 45 | 0 |
| 13l | Ipsilateral | 58 | 0 | 0 | 4 | 0 | 4 | 50 |
| 36c | Ipsilateral | 57 | 0 | 0 | 0 | 0 | 3 | 54 |
| R | Contralateral | 57 | 3 | 6 | 39 | 5 | 3 | 1 |
| TGsts | Ipsilateral | 54 | 1 | 5 | 19 | 0 | 5 | 24 |
| R | Ipsilateral | 54 | 3 | 8 | 32 | 2 | 7 | 2 |
| F6 | Ipsilateral | 54 | 0 | 0 | 10 | 0 | 11 | 33 |
| 8Bd | Ipsilateral | 53 | 0 | 0 | 2 | 1 | 7 | 43 |
| 44 | Ipsilateral | 53 | 0 | 3 | 37 | 0 | 8 | 5 |
| EC | Contralateral | 51 | 0 | 0 | 0 | 0 | 0 | 51 |
| 12o | Ipsilateral | 50 | 0 | 0 | 7 | 2 | 16 | 25 |
| 24c | Ipsilateral | 49 | 0 | 0 | 11 | 1 | 20 | 17 |
| 23b | Contralateral | 47 | 0 | 0 | 14 | 2 | 7 | 24 |
| TPO | Contralateral | 45 | 0 | 1 | 26 | 7 | 6 | 5 |
| 46f | Ipsilateral | 42 | 0 | 1 | 20 | 9 | 6 | 6 |
| Id | Contralateral | 42 | 0 | 1 | 31 | 5 | 5 | 0 |
| 7op | Ipsilateral | 41 | 3 | 2 | 12 | 2 | 7 | 15 |
| 29 | Ipsilateral | 39 | 0 | 1 | 3 | 0 | 10 | 25 |
| V6Ad | Ipsilateral | 39 | 0 | 1 | 2 | 1 | 6 | 29 |
| Iai | Ipsilateral | 37 | 0 | 0 | 0 | 0 | 0 | 37 |
| AL | Ipsilateral | 36 | 1 | 0 | 7 | 8 | 11 | 9 |
| 7b_(PFG/PF) | Ipsilateral | 31 | 1 | 1 | 6 | 4 | 5 | 14 |
| PrCO | Ipsilateral | 31 | 0 | 5 | 10 | 0 | 9 | 7 |
| LIPd | Contralateral | 31 | 0 | 6 | 10 | 1 | 6 | 8 |
| V2? | Ipsilateral | 31 | 0 | 0 | 1 | 5 | 7 | 18 |
| 5_(PEc) | Ipsilateral | 30 | 0 | 0 | 0 | 0 | 0 | 30 |
| G | Ipsilateral | 29 | 0 | 0 | 8 | 4 | 12 | 5 |
| 12l | Ipsilateral | 29 | 0 | 0 | 3 | 5 | 7 | 14 |
| A1 | Ipsilateral | 28 | 0 | 0 | 2 | 1 | 14 | 11 |
| CM | Contralateral | 25 | 5 | 7 | 12 | 1 | 0 | 0 |
| ELc | Ipsilateral | 25 | 0 | 0 | 0 | 0 | 9 | 16 |
| 24c' | Ipsilateral | 25 | 0 | 0 | 12 | 0 | 4 | 9 |
| TEad | Ipsilateral | 24 | 0 | 0 | 2 | 1 | 2 | 19 |
| 12m | Ipsilateral | 24 | 0 | 0 | 4 | 1 | 8 | 11 |
| Ig | Contralateral | 22 | 0 | 0 | 12 | 6 | 4 | 0 |
| PGa | Contralateral | 22 | 1 | 1 | 9 | 5 | 5 | 1 |
| TGdd | Ipsilateral | 21 | 0 | 1 | 5 | 0 | 9 | 6 |
| V6 | Ipsilateral | 21 | 0 | 0 | 2 | 0 | 6 | 13 |
| TGa | Ipsilateral | 20 | 0 | 0 | 1 | 0 | 4 | 15 |
| TAa | Contralateral | 19 | 1 | 1 | 8 | 3 | 4 | 2 |
| AL | Contralateral | 17 | 0 | 0 | 10 | 3 | 4 | 0 |
| 24b | Contralateral | 17 | 0 | 0 | 2 | 0 | 10 | 5 |
| CPB | Contralateral | 15 | 0 | 0 | 2 | 0 | 11 | 2 |
| ML | Ipsilateral | 15 | 0 | 0 | 4 | 2 | 5 | 4 |
| RPB | Contralateral | 14 | 0 | 0 | 0 | 3 | 11 | 0 |
| AIP | Ipsilateral | 14 | 0 | 0 | 3 | 0 | 9 | 2 |
| TH | Ipsilateral | 14 | 0 | 0 | 0 | 0 | 13 | 1 |
| Ri | Ipsilateral | 14 | 0 | 1 | 5 | 5 | 1 | 2 |
| 36r | Ipsilateral | 13 | 0 | 1 | 3 | 0 | 3 | 6 |
| IPa | Contralateral | 13 | 0 | 0 | 5 | 2 | 3 | 3 |
| 23a | Contralateral | 12 | 0 | 0 | 2 | 0 | 2 | 8 |
| 12r | Ipsilateral | 12 | 0 | 0 | 1 | 0 | 1 | 10 |
| 24b' | Contralateral | 11 | 0 | 0 | 2 | 0 | 3 | 6 |
| CL | Ipsilateral | 10 | 0 | 0 | 2 | 1 | 2 | 5 |
| ELr | Ipsilateral | 10 | 0 | 0 | 0 | 0 | 2 | 8 |
| 24c | Contralateral | 10 | 0 | 0 | 1 | 0 | 5 | 4 |
| STGr | Contralateral | 10 | 0 | 0 | 0 | 1 | 8 | 1 |
| RTp | Ipsilateral | 10 | 0 | 0 | 2 | 1 | 5 | 2 |
| LIPv | Contralateral | 9 | 0 | 0 | 4 | 0 | 3 | 2 |
| EI | Ipsilateral | 9 | 0 | 0 | 0 | 0 | 2 | 7 |
| Ial | Ipsilateral | 9 | 0 | 0 | 1 | 0 | 0 | 8 |
| MST | Contralateral | 9 | 0 | 0 | 7 | 2 | 0 | 0 |
| 30 | Contralateral | 9 | 0 | 0 | 5 | 0 | 2 | 2 |
| 8Bm | Ipsilateral | 8 | 0 | 0 | 0 | 0 | 4 | 4 |
| v23b | Contralateral | 8 | 0 | 0 | 1 | 0 | 4 | 3 |
| 35 | Contralateral | 8 | 0 | 0 | 0 | 0 | 0 | 8 |
| 8Bs | Contralateral | 7 | 0 | 0 | 3 | 0 | 2 | 2 |
| F3 | Ipsilateral | 7 | 0 | 0 | 4 | 0 | 2 | 1 |
| A1 | Contralateral | 7 | 0 | 0 | 4 | 1 | 2 | 0 |
| RT | Ipsilateral | 7 | 0 | 2 | 4 | 0 | 1 | 0 |
| 5_(PEa) | Ipsilateral | 7 | 0 | 0 | 0 | 0 | 0 | 7 |
| Iapl | Ipsilateral | 7 | 0 | 0 | 0 | 0 | 0 | 7 |
| 1/2 | Ipsilateral | 6 | 0 | 0 | 0 | 0 | 3 | 3 |
| F4 | Ipsilateral | 6 | 0 | 1 | 2 | 0 | 2 | 1 |
| TEa | Contralateral | 6 | 0 | 0 | 3 | 1 | 2 | 0 |
| 24a' | Ipsilateral | 5 | 0 | 1 | 2 | 0 | 2 | 0 |
| V1 | Ipsilateral | 5 | 0 | 0 | 1 | 0 | 2 | 2 |
| 9m | Ipsilateral | 5 | 0 | 0 | 0 | 0 | 2 | 3 |
| RTL | Ipsilateral | 5 | 0 | 0 | 0 | 1 | 3 | 1 |
| 24a | Ipsilateral | 5 | 0 | 0 | 3 | 0 | 2 | 0 |
| 24a | Contralateral | 5 | 0 | 0 | 1 | 0 | 2 | 2 |
| 24c' | Contralateral | 5 | 0 | 0 | 2 | 0 | 0 | 3 |
| Tpt | Contralateral | 4 | 0 | 0 | 0 | 0 | 0 | 4 |
| ECL | Ipsilateral | 4 | 0 | 0 | 0 | 0 | 0 | 4 |
| SII | Contralateral | 4 | 0 | 0 | 2 | 1 | 1 | 0 |
| TGdg | Ipsilateral | 4 | 0 | 0 | 0 | 0 | 1 | 3 |
| F2_(6DR/6DC) | Contralateral | 4 | 0 | 0 | 0 | 0 | 2 | 2 |
| Iam | Ipsilateral | 3 | 0 | 0 | 0 | 0 | 0 | 3 |
| 29 | Contralateral | 3 | 0 | 0 | 0 | 0 | 1 | 2 |
| V4 | Contralateral | 3 | 0 | 0 | 1 | 0 | 2 | 0 |
| 31 | Contralateral | 3 | 0 | 0 | 0 | 0 | 3 | 0 |
| 8Bd | Contralateral | 3 | 0 | 0 | 1 | 0 | 0 | 2 |
| TGvd | Ipsilateral | 3 | 0 | 0 | 0 | 0 | 2 | 1 |
| F1_(4) | Ipsilateral | 3 | 0 | 0 | 1 | 2 | 0 | 0 |
| 23c | Contralateral | 3 | 0 | 0 | 1 | 0 | 0 | 2 |
| Pi | Contralateral | 3 | 0 | 1 | 2 | 0 | 0 | 0 |
| V2_or_v23b? | Contralateral | 3 | 0 | 0 | 1 | 0 | 1 | 1 |
| 11l | Ipsilateral | 2 | 0 | 0 | 0 | 0 | 1 | 1 |
| 46d | Contralateral | 2 | 0 | 0 | 1 | 0 | 1 | 0 |
| 24a' | Contralateral | 2 | 0 | 0 | 1 | 0 | 1 | 0 |
| ER | Ipsilateral | 2 | 0 | 0 | 0 | 0 | 1 | 1 |
| 9d | Contralateral | 2 | 0 | 0 | 1 | 0 | 0 | 1 |
| ECL | Contralateral | 2 | 0 | 0 | 0 | 0 | 0 | 2 |
| 7b_(PFG/PF) | Contralateral | 2 | 0 | 1 | 1 | 0 | 0 | 0 |
| TEm | Contralateral | 2 | 0 | 0 | 0 | 0 | 2 | 0 |
| F6 | Contralateral | 2 | 0 | 0 | 0 | 0 | 0 | 2 |
| 36p | Ipsilateral | 2 | 0 | 0 | 0 | 0 | 0 | 2 |
| TEpv | Contralateral | 2 | 0 | 0 | 0 | 0 | 1 | 1 |
| TF | Contralateral | 2 | 0 | 0 | 0 | 0 | 1 | 1 |
| TH | Contralateral | 2 | 0 | 0 | 0 | 0 | 2 | 0 |
| Ia | Ipsilateral | 1 | 0 | 0 | 0 | 0 | 0 | 1 |
| VIP | Contralateral | 1 | 0 | 0 | 0 | 0 | 0 | 1 |
| 7op | Contralateral | 1 | 0 | 0 | 1 | 0 | 0 | 0 |
| 5_(PE) | Contralateral | 1 | 0 | 0 | 0 | 0 | 1 | 0 |
| TEav | Contralateral | 1 | 0 | 0 | 1 | 0 | 0 | 0 |
| 32 | Ipsilateral | 1 | 0 | 0 | 0 | 0 | 1 | 0 |
| 10mr | Ipsilateral | 1 | 0 | 0 | 0 | 0 | 0 | 1 |
| 1/2 | Contralateral | 1 | 0 | 0 | 0 | 1 | 0 | 0 |
| TFO | Contralateral | 1 | 0 | 0 | 0 | 0 | 1 | 0 |
| 46v | Contralateral | 1 | 0 | 0 | 1 | 0 | 0 | 0 |
| LOP | Contralateral | 1 | 0 | 0 | 0 | 0 | 1 | 0 |
| 11m | Ipsilateral | 1 | 0 | 0 | 0 | 1 | 0 | 0 |
| 45a | Contralateral | 1 | 0 | 0 | 1 | 0 | 0 | 0 |
| F7 | Contralateral | 1 | 0 | 0 | 0 | 0 | 1 | 0 |
| F5_(6Va/6Vb) | Contralateral | 1 | 0 | 0 | 1 | 0 | 0 | 0 |
| V2 | Contralateral | 1 | 0 | 0 | 0 | 0 | 0 | 1 |
| TGsts | Contralateral | 1 | 0 | 0 | 1 | 0 | 0 | 0 |
| 8Ad | Contralateral | 1 | 0 | 0 | 0 | 1 | 0 | 0 |
| TGvg | Ipsilateral | 1 | 0 | 0 | 0 | 0 | 0 | 1 |
| Ri | Contralateral | 1 | 0 | 0 | 1 | 0 | 0 | 0 |

**Table S9. Index of abbreviations of M132 parcellation**

| **Abbreviation** | **Fullname** |
| --- | --- |
| 10 | frontopolar cortex (area 10) |
| 11 | orbital prefrontal cortex (area 11) |
| 12 | orbital prefrontal cortex (area 12) |
| 13 | orbital prefrontal cortex (area 13) |
| 14 | orbital prefrontal cortex (area 14) |
| 25 | medial prefrontal area (subgenual cortex) |
| 44 | ventro-lateral prefrontal area in the arcuate sulcus lower limb |
| 45A | ventro-lateral prefrontal area |
| 45B | ventro-lateral prefrontal area in the arcuate sulcus lower limb |
| 46d | dorso-lateral prefrontal area in the dorsal bank and lip of the principal sulcus |
| 46v | ventro-lateral prefrontal area in the ventral bank and lip of the principal sulcus |
| 8B | dorso-lateral / dorso-medial prefrontal area |
| 8l | area 8, lateral subdivision |
| 8m | area 8, medial subdivision |
| 8r | area 8, rostral subdivision |
| 9 | dorso-lateral prefrontal area |
| 9/46d | dorso-lateral prefrontal area 9/46, dorsal |
| 9/46v | dorso-lateral prefrontal area 9/46, ventral |
| PIR | piriform cortex |
| ProM | proisocortex motor |
| F1 | agranular frontal area F1 (or 4) |
| F2 | agranular frontal area F2 (or 6DR/6DC) |
| F3 | agranular frontal area F3 (or SMA) |
| F4 | agranular frontal area F4 (or 4C/6Va/6Vb) |
| F5 | agranular frontal area F5 (or 6Va/6Vb) |
| F6 | agranular frontal area F6 (or preSMA) |
| F7 | agranular frontal area F7 (or 6DR) |
| INS | insular cortex |
| 1 | primary somatosensory area 1 |
| 2 | primary somatosensory area 2 |
| 3 | primary somatosensory area 3 |
| Gu | gustatory cortex |
| SII | secondary somatosensory area |
| PERI | perirhinal cortex |
| ENTO | entorhinal cortex |
| FST | floor of superior temporal area |
| IPa | area IPa (sts fundus) |
| MST | medial superior temporal area |
| MTc | middle temporal cortex |
| PGa | sts fundus/dorsal bank area |
| Pro.St. | prostriate cortex |
| POLE | temporal pole |
| STPc | superior temporal polysensory area, caudal |
| STPi | superior temporal polysensory area, intermediate |
| STPr | superior temporal polysensory area, rostral |
| TEO | area TEO |
| TEOm | temporal-occipital area, medial |
| TEa/ma | temporal area TEa/ma |
| TEa/mp | temporal area TEa/mp |
| TEad | dorsal subregion of anterior TE |
| TEav | ventral subregion of anterior TE |
| TEpd | dorsal subregion of posterior TE |
| TEpv | ventral subregion of posterior TE |
| TF/TH | parahippocampal areas TF/TH |
| TPt | sts dorsal bank area (temporoparietal) |
| Pi | inferior pulvinar |
| 5 | superior parietal lobule area (or PE/PEa/PEc) |
| 7A | caudal inferior parietal lobule area (or Opt/PG) |
| 7B | rostral inferior parietal lobule area (or PFG/PF) |
| 7m | medial parietal area (or PGm) |
| 7op | parietal opercular area |
| AIP | anterior intraparietal area |
| DP | dorsal prelunate area |
| LIP | lateral intraparietal area |
| MIP | medial intraparietal area |
| OPAI | orbital periallocortex |
| OPRO | orbital proisocortex |
| PIP | posterior intraparietal area |
| V6 | visual area 6 (or PO) |
| V6A | visual area 6A |
| VIP | ventral intraparietal area |
| Core | core region of auditory cortex (AI/R/RT) |
| LB | lateral belt of auditory cortex |
| MB | medial belt of auditory cortex |
| PBc | caudal parabelt region of auditory cortex |
| PBr | rostral parabelt region of auditory cortex |
| 23 | posterior cingulate cortex (area 23) |
| 24a | subregion of anterior cingulate cortex |
| 24b | subregion of anterior cingulate cortex |
| 24c | subregion of anterior cingulate cortex |
| 24d | anterior cingulate cortex (area 24d) |
| 29/30 | retrosplenial cortex (areas 29 and 30) |
| 31 | posterior cingulate area |
| 32 | medial prefrontal area |
| MTp | middle temporal area, posterior |
| V1c | primary visual cortex, central representation |
| V1fpLF | V1 foveal periphery, lower field |
| V1fpUF | V1 foveal periphery, upper field |
| V1pcLF | V1 peripheral, lower field |
| V1pcUF | V1 peripheral, upper field |
| V2c | visual area 2, central |
| V2fpLF | V2 foveal periphery, lower field |
| V2fpUF | V2 foveal periphery, upper field |
| V2pcLF | V2 peripheral, lower field |
| V2pcUF | V2 peripheral, upper field |
| V3A | visual area V3A |
| V3c | visual area 3, central |
| V3fpLF | V3 foveal periphery, lower field |
| V3fpUF | V3 foveal periphery, upper field |
| V3pcLF | V3 peripheral, lower field |
| V3pcUF | V3 peripheral, upper field |
| V4LF | visual area 4, lower field |
| V4UF | visual area 4, upper field |
| V4c | visual area 4, central |
| V4t | V4 transitional area |

**Table S10. Index of abbreviations of SARM parcellations**

| **Abbreviation** | **Fullname** |
| --- | --- |
| Cl | claustrum |
| DEn | dorsal_endopiriform_claustrum |
| VEn | ventral_endopiriform_claustrum |
| Pir | piriform_cortex |
| Hi | hippocampus |
| S | subicular_complex |
| fi | fimbria |
| f | fornix |
| BM | basomedial_amygdaloid_nucleus |
| VCo | ventral_cortical_amygdaloid_nucleus |
| AHi | amygdalohippocampal_area |
| LaD | lateral_dorsal_amygdaloid_nucleus |
| LaV | lateral_ventral_amygdaloid_nucleus |
| APir | amygdalopiriform_transition_area |
| BLD | basolateral_dorsal_amygdaloid_nucleus |
| BLI | basolateral_intermediate_amygdaloid_nucleus |
| BLV | basolateral_ventral_amygdaloid_nucleus |
| PaL | paralaminar_amygdaloid_nucleus |
| Ce | central_amygdaloid_nucleus |
| Me | medial_amygdaloid_nucleus |
| AA | anterior_amygdaloid_area |
| EA | extended_amygdala |
| IPAC | posterior_interstitial_nucleus |
| STIA | intraamygdaloid_stria_terminalis |
| CdH | caudate_head |
| CdT | caudate_tail |
| Pu | putamen |
| ASt | amygdalostriatal_transition |
| ic | internal_capsule |
| Acb | accumbens |
| Tu | olfactory_tubercule |
| EGP | external_globus_pallidus |
| IGP | internal_globus_pallidus |
| al | ansa_lenticularis |
| ac | anterior_commissure |
| VP | ventral_pallidum |
| PeB | peri-basal_region |
| B | basal_nucleus |
| ST | bed_nucleus_of_the_stria_terminalis |
| SFi | septimbrial_nucleus |
| SDB | septum_diagonal_band |
| opt | optic_tract |
| PO | preoptic_area |
| LTer-VOLT | lamina_terminalis |
| Pa | paraventricular_hypothalamus |
| VMH | ventromedial_hypothalamic_nucleus |
| Arc | arcuate_nucleus |
| SOpt | supraoptic_hypothalamus |
| AH | anterior_hypothalamic_area |
| DM | dorsal_medial_hypothalamic_nucleus |
| LH | lateral_hypothalamic_area |
| JPLH | justaparaventricular_lateral_hypothalamus |
| PLH | peduncular_lateral_hypothalamus |
| STh | subthalamic_nucleus |
| PH | posterior_hypothalamic_nucleus |
| PeF | prefornical_hypothalamus |
| RM | retromammillary_hypothalamus |
| MM | mammillary_hypothalamus |
| ZI-H | zona_incerta_and_lenticular_fascicles |
| Rt | reticular_thalamus |
| A | anterior_thalamus |
| cap | capsule_of_the_antherior_thalamus |
| CL-PC | centrolateral_and_paracentral_thalamus |
| CMn-PF | centromedial-parafascicular_thalamus |
| SPFPC | subparafascicular_nucleus |
| SG | supra_geniculate |
| Lim | limitans_thalamus |
| MD | mediodorsal_thalamus |
| IMD | intermediodorsal_thalamus |
| PT-PV-sm | paratenial-paraventral-thalamus |
| PVP | paraventricular_posterior_thalamus |
| PVA | paraventricular_anterior_thalamus |
| IAM | interanteromedial_thalamus |
| CM | central_medial_thalamus |
| Re-Rh-Xi | Reuniens-rhomboid-xiphoid_thalamus |
| VA | ventral_anterior_thalamus |
| VLA | ventral_lateral_anterior_thalamus |
| VLX | ventral_lateral_x_thalamus |
| VLPV | ventral_lateral_posteroventral_thalamus |
| VLPD | ventral_lateral_posterodorsal_thalamus |
| VPM-VPL | ventroposterior_medial_and_lateral_thalamus |
| VMPo-VMB | ventromedial_posterior+basal_thalamus |
| VPI | ventroposterior_inferior_thalamus |
| VM | ventral_medial_thalamus |
| LP | lateral_posterior_thalamus |
| LD | lateral_dorsal_thalamus |
| APul | anterior_pulvinar |
| MPul | medial_pulvinar |
| LPul | lateral_pulvinar |
| IPul | inferior_pulvinar |
| bsc | brachium_superior_colliculus |
| DLG | dorsal_lateral_geniculate |
| MG | medial_geniculate |
| Pi | pineal_gland |
| Hb | habenula |
| pc | posterior_commissure |
| PrC | precommissural_nucleus |
| PCom_MCPC | posterior_commissural_nuclei |
| p1Rt | p1_mesencephalic_reticulum |
| SC | superior_colliculus |
| ICo | inferior_colliculus |
| Sag-RL | saginum_nucleus |
| n4 | fourth_nerve |
| PAG | periaqueductal_gray |
| PBG | parabigeminal_nucleus |
| MiTg | mircrocellular_tegmentum |
| PrCnF | precuneiform_nucleus |
| CnF | cuneiform_nucleus |
| VLTg | ventrolateral_tegmentum |
| PTg | pedunculopontine_nucleus |
| ATg | anterior_tegmental_nucleus |
| mRt | mesencephalic_reticular_formation |
| RIs | retroisthmic_nucleus |
| OM | occulomotor_nuclei |
| mlf | mediolateral_fascicle |
| MR | medial_Raphe |
| DR | dorsal_Raphe |
| LiR | linear_Raphe |
| R | red_nucleus |
| PR-RI | prerubral_field |
| RF | retrorubral_field |
| SN | substantia_nigra |
| VTA | ventral_tegmental_area |
| IP | interpeduncular_nucleus |
| n3 | occulomotor_nerve |
| cp | cerebral_peduncle |
| LPB | lateral_parabrachial_nucleus |
| MPB | medial_parabrachial_nucleus |
| scp | superior_cerebellar_peduncle |
| KF | Kolliker-Fuse_nucleus |
| LC-Me5 | locus-coeruleus_region |
| me5 | mesencephalic_trigeminal_nerve |
| CG | central_gray_nucleus |
| LDTg | lateral_dorsal_tegmentum |
| MDTg | medial_dorsal_tegmentum |
| ll | lateral_lemniscus |
| DLL | dorsal_lateral_leminscus_nucleus |
| ILL-VLL | intermediate+ventral_lateral_lemniscus_nuclei |
| SubC | subceruleus |
| RtTg | reticulotegmentum |
| xscp | superior_cerebellar_peduncle_decussation |
| PnO | pontine_reticulum_oral |
| PnC | pontine_reticulum_caudal |
| SO | superior_olive |
| ml | medial_lemniscus |
| tz | trapezoid_bundle_region |
| Pn | pontine_nucleus |
| s5 | sensory_root_of_the_trigeminal_nerve |
| tfp | transverse_fibers |
| mcp | medial_cerebellar_peduncle |
| icp | inferior_cerebellar_peduncle |
| oc | olivocerebellar_tract |
| Lat | lateral_cerebellar_nucleus |
| IntA | anterior_interposed_cerebellar_nucleus |
| IntP | posterior_interposed_cerebellar_nucleus |
| Fas | fastigial_cerebellar_nucleus |
| Fl | flocculus |
| PFl | paraflocculus |
| Cb1V | cerebellar_lobule_1_vermis |
| Cb2V | cerebellar_lobule_2_vermis |
| Cb3V | cerebellar_lobule_3_vermis |
| Cb4V | cerebellar_lobule_4_vermis |
| Cb5V | cerebellar_lobule_5_vermis |
| Cb6V | cerebellar_lobule_6_vermis |
| Cb7V | cerebellar_lobule_7_vermis |
| Cb8V | cerebellar_lobule_8_vermis |
| Cb9V | cerebellar_lobule_9_vermis |
| Cb10V | cerebellar_lobule_10_vermis |
| Cb3I | cerebellar_lobule_3_intermediate |
| Cb4I | cerebellar_lobule_4_intermediate |
| Cb5I | cerebellar_lobule_5_intermediate |
| Cb6I | cerebellar_lobule_6_intermediate |
| Cop | copula_of_the_pyramis |
| Sim | simple_lobule |
| PM | paramedian_lobule |
| Crus1 | ansiform_lobule_crus1 |
| Crus2 | ansiform_lobule_crus2 |
| Ve | vestibular_nucleus |
| DVC | cochlear_nucleus |
| n8 | vestibulochoclear_nerve |
| Pr | prepositus_nucleus |
| Nu5 | motor_trigeminal_nucleus |
| Pr5 | principal_trigeminal_nucleus |
| Sp5O | spinal_trigeminal_nucleus_oral |
| Sp5I | spinal_trigeminal_nucleus_intermediate |
| Sp5C | spinal_trigeminal_nucleus_caudal |
| CuGr | cuneate_gracile_nuclei |
| Sol | solitary_tract_nucleus |
| Nu10-12 | visceral_motor_nuclei |
| Nu6 | abducens_motor_nucleus |
| Nu7 | facial_motor_nucleus |
| PCIRt | parvicellular_and_intermediate_reticular_nuclei |
| DPGi | dorsal_paragigantocellular_reticular_nucleus |
| Gi | gigantocellular_reticular_nucleus |
| MdRt | medullar_reticular_nucleus |
| RMg | Raphe_magnus_nucleus |
| ROb | Raphe_obscurus_nucleus |
| IO | inferior_olive |
| LRt | lateral_reticular_nucleus |
| py | pyramidal_tract |

**Table S11. Index of abbreviations of D99 parcellations**

| **Abbreviation** | **Fullname** |
| --- | --- |
| 8Bm | dorso-lateral prefrontal area 8Bm |
| 11l | orbital prefrontal area 11l |
| 14r | orbito-medial prefrontal area 14r (gyrus rectus) |
| Ia | agranular insula (Ia) |
| 13b | orbital prefrontal area 13b |
| 36c | area 36 of the perirhinal cortex, caudal subregion |
| 24a' | subregion 24a' of anterior cingulate cortex |
| 10mc | dorso-medial prefrontal area 10mc |
| CPB | caudal parabelt region (CPB) of the auditory cortex |
| 29 | retrosplenial cortex area 29 |
| v23b | subregion v23b of posterior cingulate cortex |
| 13l | orbital prefrontal area 13l |
| V4 | visual area 4, dorsal part (V4) |
| 31 | posterior cingulate area 31 |
| Ig | granular insula (Ig) |
| ML | middle lateral belt region (ML) of the auditory cortex |
| 23a | subregion 23a of posterior cingulate cortex |
| ER | entorhinal cortex, rostral division (ER) |
| 10o | medial prefrontal area 10o |
| VIP | ventral intraparietal area (VIP) |
| LIPv | lateral intraparietal area, ventral subdivision (LIPv) |
| 8Bs | dorso-medial prefrontal area 8Bs |
| V1 | visual area 1 (primary visual cortex, V1) |
| G | gustatory cortex (G) |
| 9d | dorso-lateral prefrontal area 9d in the arcuate sulcus upper limb |
| 7op | parietal opercular area 7op |
| V4t | V4 transitional area (V4t) |
| 46f | ventro-lateral prefrontal area 46f in the ventral bank and lip of the principal sulcus (ps) |
| 5_(PE) | superior parietal lobule area 5 (PE) |
| TEav | ventral subregion of anterior TE (TEav) |
| 32 | medial prefrontal area 32 |
| AL | anterior lateral belt region (AL) of the auditory cortex |
| 10mr | medial prefrontal area 10mr |
| 24b | subregion 24b of anterior cingulate cortex |
| 1月2日 | somatosensory areas 1 and 2 |
| 8Av | periarcuate area 8Av (or frontal eye field), dorsal subdivision |
| ECL | entorhinal cortex, caudal limiting division (ECL) |
| FST | floor of superior temporal area (FST) |
| 12m | ventro-lateral prefrontal area 12m |
| V6Av | visual area 6A, ventral division (V6Av, or PO) |
| 8Bd | periarcuate area 8Bd (or frontal eye field), ventral subdivision |
| TGvd | sts part of the temporal pole (TGvd) |
| V3d | visual area 3, dorsal part (V3d) |
| F1_(4) | agranular frontal area F1 (or area 4) |
| Iam | medial agranular insula area (Iam) |
| SII | secondary somatosensory area (SII) |
| 23b | subregion 23b of posterior cingulate cortex |
| 13m | orbital prefrontal area 13m |
| 9m | dorso-lateral prefrontal area 9m |
| EI | entorhinal cortex, intermediate division (EI) |
| RPB | rostral parabelt region (RPB) of the auditory cortex |
| 13a | ventro-lateral prefrontal area 13a |
| F3 | agranular frontal area F3 (or SMA) |
| R | reticular nucleus / rostral core region (R) of the auditory cortex |
| V6 | visual area 6 (V6, or PO) |
| TFO | area TFO of the parahippocampal cortex |
| V4v | visual area 4, ventral part (V4v) |
| 46v | ventro-lateral prefrontal area 46v in the ventral bank and lip of the principal sulcus (ps) |
| 5_(PEc) | superior parietal lobule area 5 (PEc) |
| TGdd | dorsal temporal pole (TGdd) |
| 35 | area 35 of the perirhinal cortex |
| A1 | auditory area A1, core region of the auditory cortex |
| Ial | lateral agranular insula area (Ial) |
| 7m_(PGm) | medial parietal area 7m (PGm) |
| v23b? | subregion v23b of posterior cingulate cortex (uncertain) |
| 23c | subregion 23c of posterior cingulate cortex |
| EC | entorhinal cortex, caudal division (EC) |
| Id | dysgranular insula (Id) |
| MIP | medial intraparietal area (MIP) |
| PIP | posterior intraparietal area (PIP) |
| 7b_(PFG/PF) | rostral inferior parietal lobule area 7b (PFG/PF) |
| TEpd | dorsal subregion of posterior TE (TEpd) |
| V3v | visual area 3, ventral part (V3v) |
| 44 | ventro-lateral prefrontal area 44 in the arcuate sulcus lower limb (asl) |
| MT | middle temporal area (MT) |
| TEm | sts ventral bank area (TEm) |
| Tpt | sts dorsal bank area (Tpt) |
| 24c | subregion 24c of anterior cingulate cortex |
| MST | medial superior temporal area (MST) |
| RM | rostromedial belt region (RM) of the auditory cortex |
| STGr | rostral superior temporal gyrus (STGr) |
| Pi | inferior pulvinar (Pi) |
| F6 | agranular frontal area F6 (or preSMA) |
| DP | dorsal prelunate area (DP) |
| 12r | orbital prefrontal area 12r |
| PrCO | precentral opercular area (PrCO) |
| PEci | superior parietal lobule area (PEci), cingulate sulcus |
| TGdg | dysgranular part of the dorsal temporal pole (TGdg) |
| AIP | anterior intraparietal area (AIP) |
| 36p | area 36 of the perirhinal cortex, temporal-polar subregion |
| LOP | lateral occipital parietal area (LOP) |
| 11m | orbital prefrontal area 11m |
| RTp | rostrotemporal area (RTp, p refers to polar) |
| 25 | medial prefrontal area (subgenual cortex) |
| Pir | piriform cortex (Pir) |
| Iapm | posteromedial agranular insula area (Iapm) |
| RT | rostrotemporal core region (RT) of the auditory cortex |
| 7a_(Opt/PG) | caudal inferior parietal lobule area 7a (Opt/PG) |
| TEpv | ventral subregion of posterior TE (TEpv) |
| V3A | visual area V3A |
| 45a | ventro-lateral prefrontal area 45a |
| TEO | area TEO |
| F7 | agranular frontal area F7 (or 6DR) |
| 46d | dorso-lateral prefrontal area 46d in the dorsal bank and lip of the principal sulcus (ps) |
| Iai | intermediate agranular insula area (Iai) |
| F5_(6Va/6Vb) | agranular frontal area F5 (or 6Va/6Vb) |
| LIPd | lateral intraparietal area, dorsal subdivision (LIPd) |
| V2 | visual area 2 (V2) |
| 36r | area 36 of the perirhinal cortex, rostral subregion |
| 12l | orbital prefrontal area 12l |
| 5_(PEa) | superior parietal lobule area 5 (PEa) |
| TGsts | granular part of the dorsal temporal pole (TGsts) |
| RTM | medial rostrotemporal belt region (RTM) of the auditory cortex |
| 3a/b | somatosensory areas 3a and 3b |
| TGa | agranular part of the temporal pole (TGa) |
| V6Ad | visual area 6A, dorsal division (V6Ad) |
| TF | area TF of the parahippocampal cortex |
| 45b | ventro-lateral prefrontal area 45b in the arcuate sulcus lower limb (asl) |
| TEad | dorsal subregion of anterior TE (TEad) |
| PGa | sts fundus/dorsal bank area (PGa) |
| F4 | agranular frontal area F4 (or 4C/6Va/6Vb) |
| 8Ad | transitional area 8Ad near the lip of ips |
| RTL | lateral rostrotemporal belt region (RTL) of the auditory cortex |
| 12o | orbital prefrontal area 12o |
| CM | caudomedial belt region (CM) of the auditory cortex |
| TAa | sts dorsal bank area (TAa) |
| F2_(6DR/6DC) | agranular frontal area F2 (or 6DR/6DC) |
| TGvg | ventral temporal pole (TGvg) |
| CL | caudal lateral belt region (CL) of the auditory cortex |
| TPO | temporo-parietal area (TPO) |
| TEa | sts ventral bank area (TEa) |
| IPa | area IPa (sts fundus) |
| ELr | entorhinal cortex, lateral division, rostral part (ELr) |
| ELc | entorhinal cortex, lateral division, caudal part (ELc) |
| V2? | visual area 2 (V2, uncertain) |
| V2_or_v23b? | visual area 2 or subregion of posterior cingulate cortex (V2 or v23b, uncertain) |
| 30 | retrosplenial cortex area 30 |
| 24b' | subregion 24b' of anterior cingulate cortex |
| 24a | subregion 24a of anterior cingulate cortex |
| 24c' | subregion 24c' of anterior cingulate cortex |
| 14c | orbital prefrontal area 14c |
| EO | entorhinal cortex, olfactory division (EO) |
| TH | ventral dysgranular part of the temporal pole (TH) |
| Pir/PACo | periamygdaloid cortex/piriform cortex (PACo/Pir) |
| Iapl | posterolateral agranular insula area (Iapl) |
| Ri | retroinsula (Ri) |

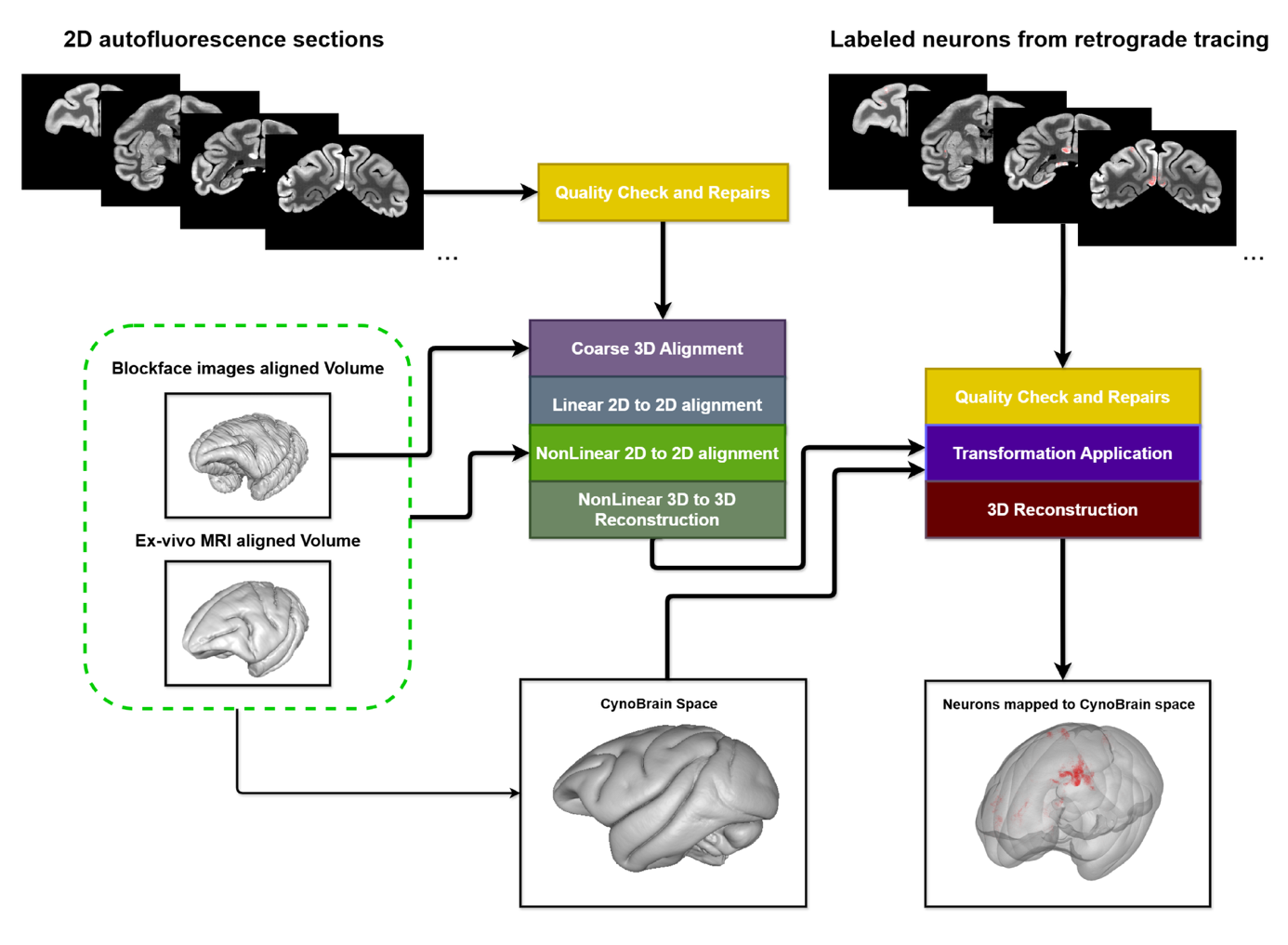
**Figure S1. Pipeline for autofluorescence section reconstruction and labled neuron mapping to CynoBrain space.**

2D autofluorescence sections were processed through hierarchical 2D-to-2D and 3D-to-3D nonlinear registration, using blockface and ex-vivo MRI volumes as references. In parallel, sections containing retrogradely labeled neurons underwent quality control, transformation application, and 3D reconstruction. Both streams converge to map labeled neurons into the CynoBrain common space

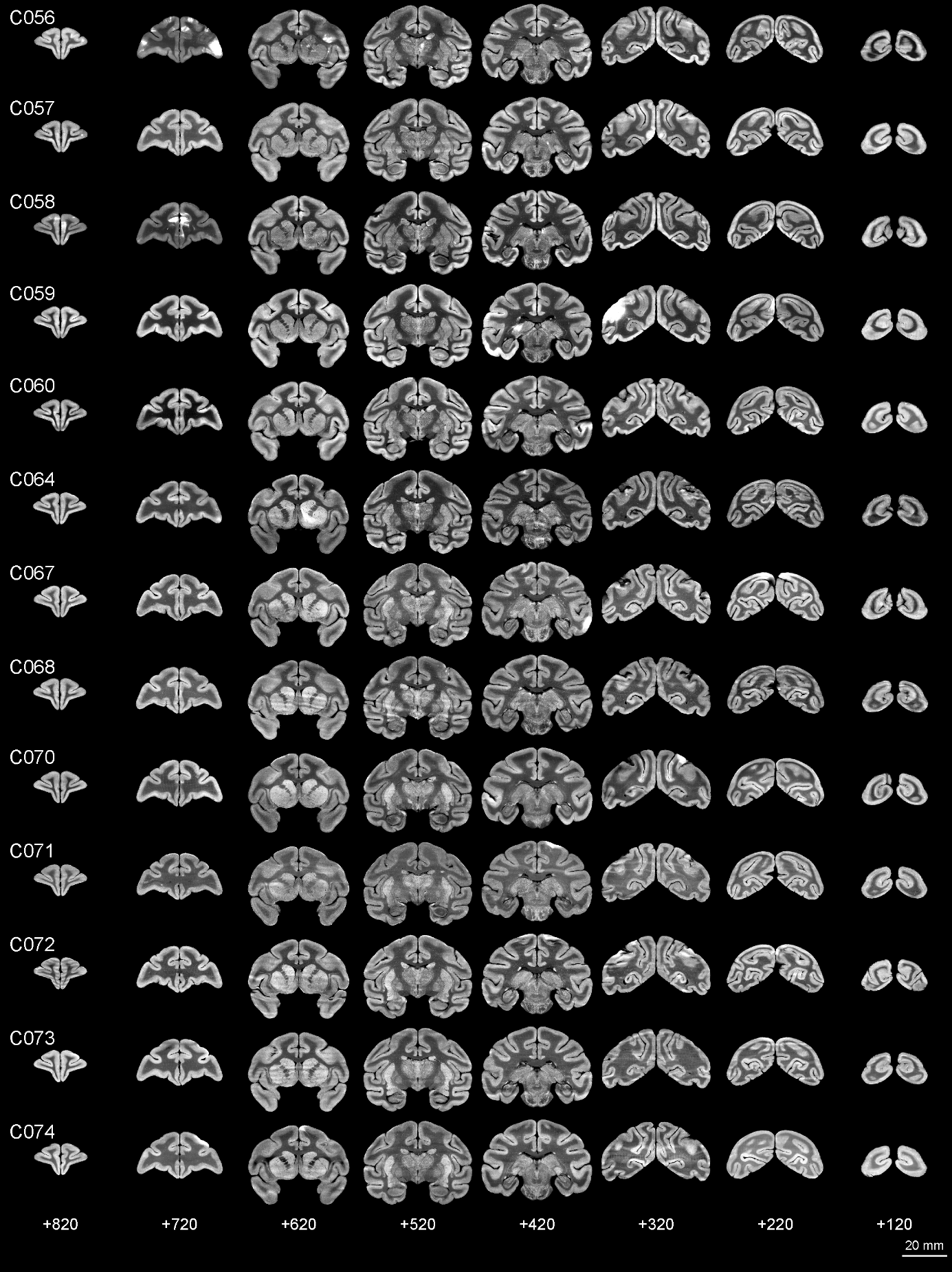

**Figure S2. Individual autofluorescence brain volumes reconstructed in CynoBrain space.**

Coronal sections from all 13 macaque subjects (C056–C074) used to construct the high-resolution autofluorescence template are shown. For each individual, 8 evenly spaced coronal slices spanning the full anterior-posterior extent of the brain are displayed, demonstrating the quality of inter-subject alignment. Images are shown in grayscale with consistent intensity scaling across subjects.

**Figure S3. 2D layer delineation derived from transcriptome atlas of macaque in native space.**

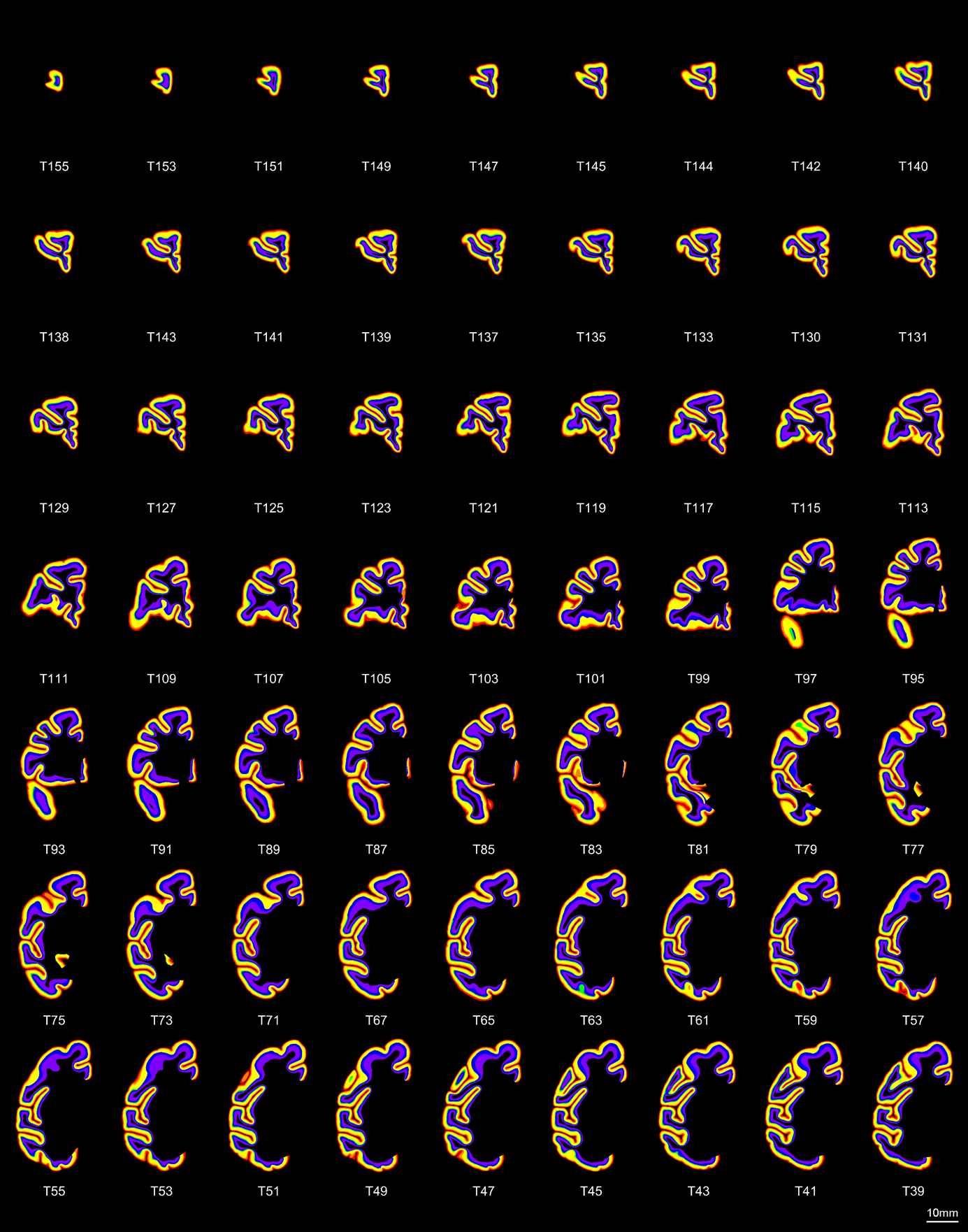

Coronal sections showing the layer segments in the macaque brain delineated in transcriptome atlas of macaque from Chen et al., Cell 2023.

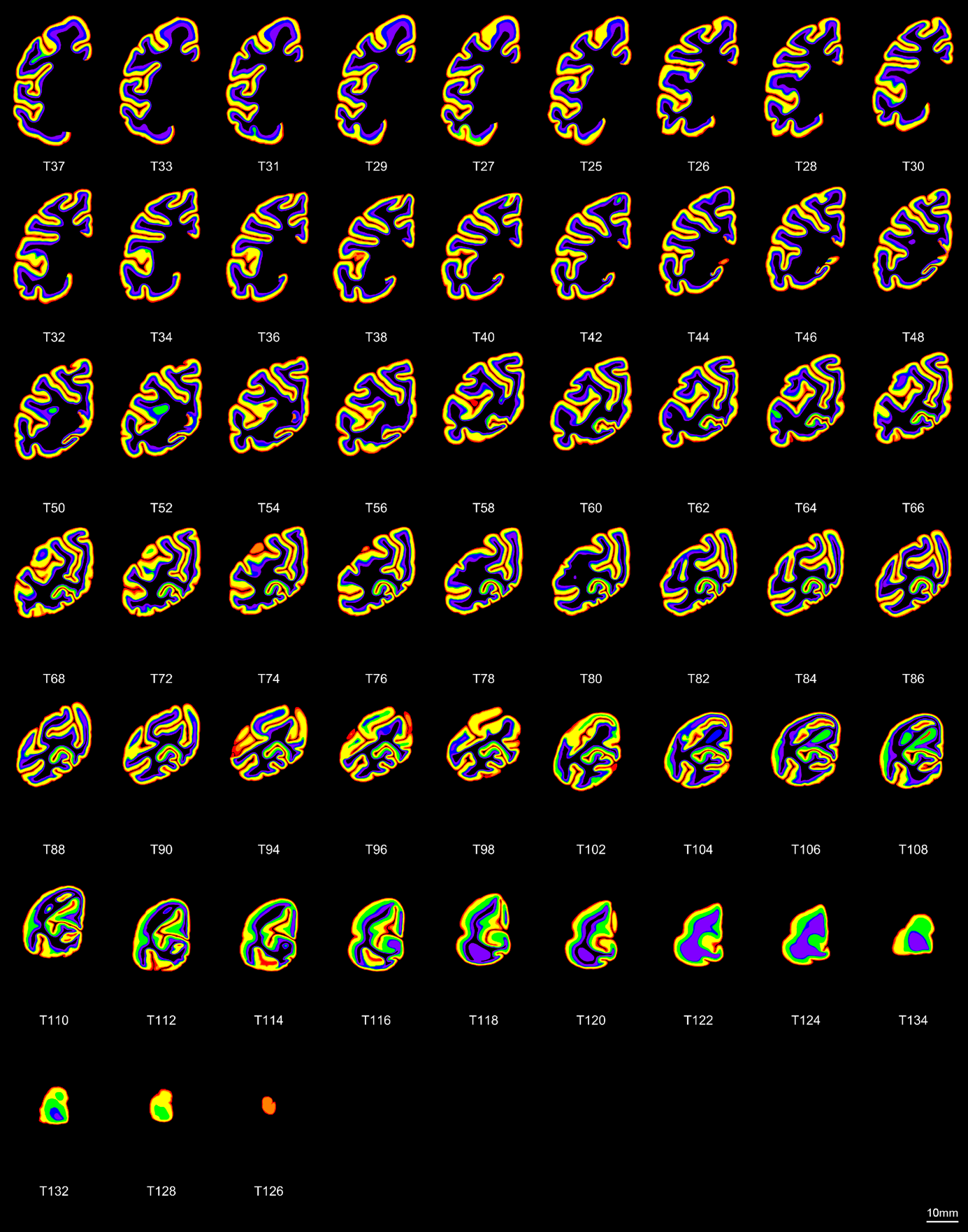

**Figure S3. 2D layer delineation derived from transcriptome atlas of macaque in native space. (Continued)**

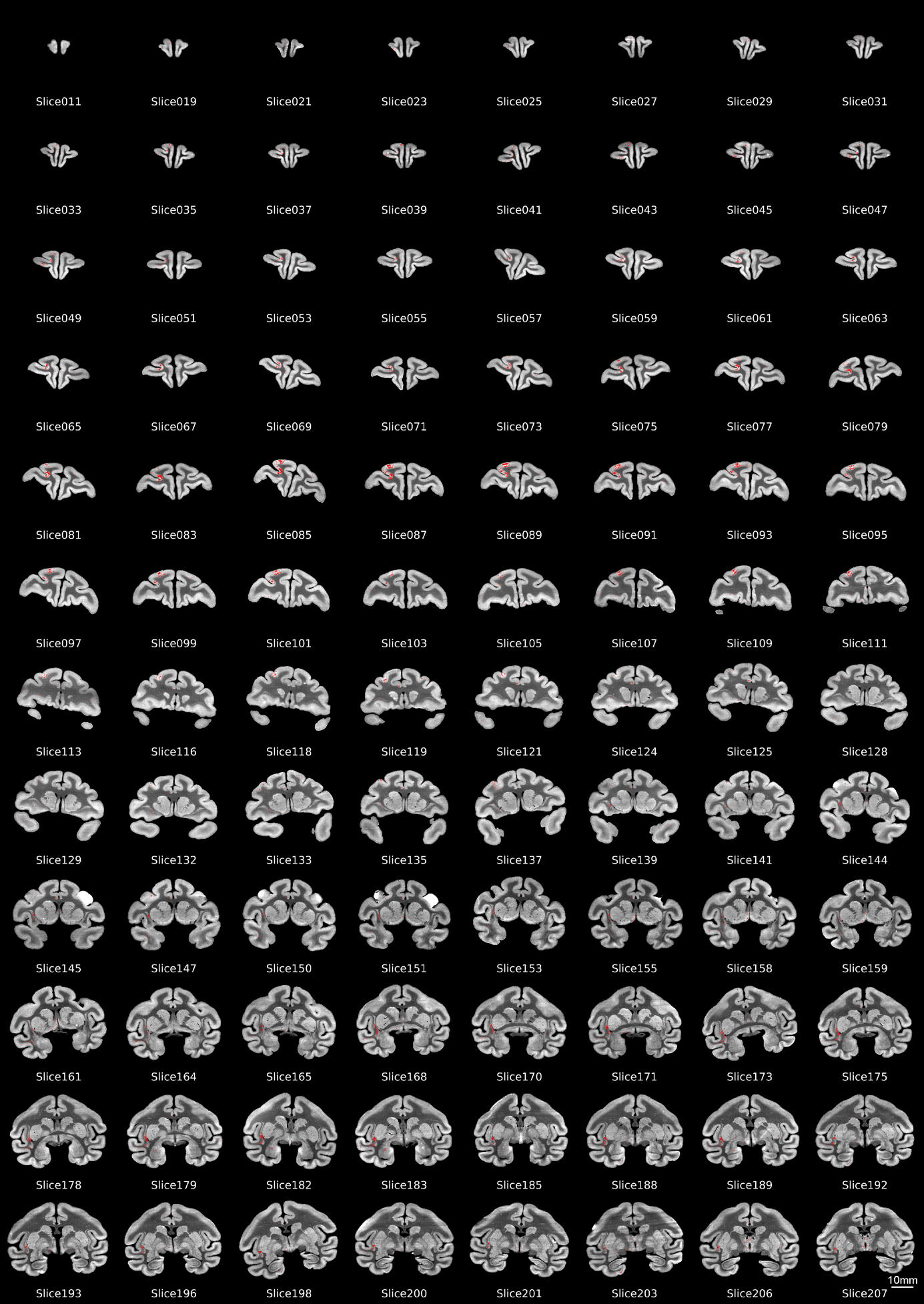

**Figure S4. Distribution of neurons projecting to area 23 in the native space.**

Coronal sections showing the distribution of retrogradely labeled neurons in the macaque brain prior to 3D reconstruction.Red dots represent retrogradely labeled neurons projecting to area 23.

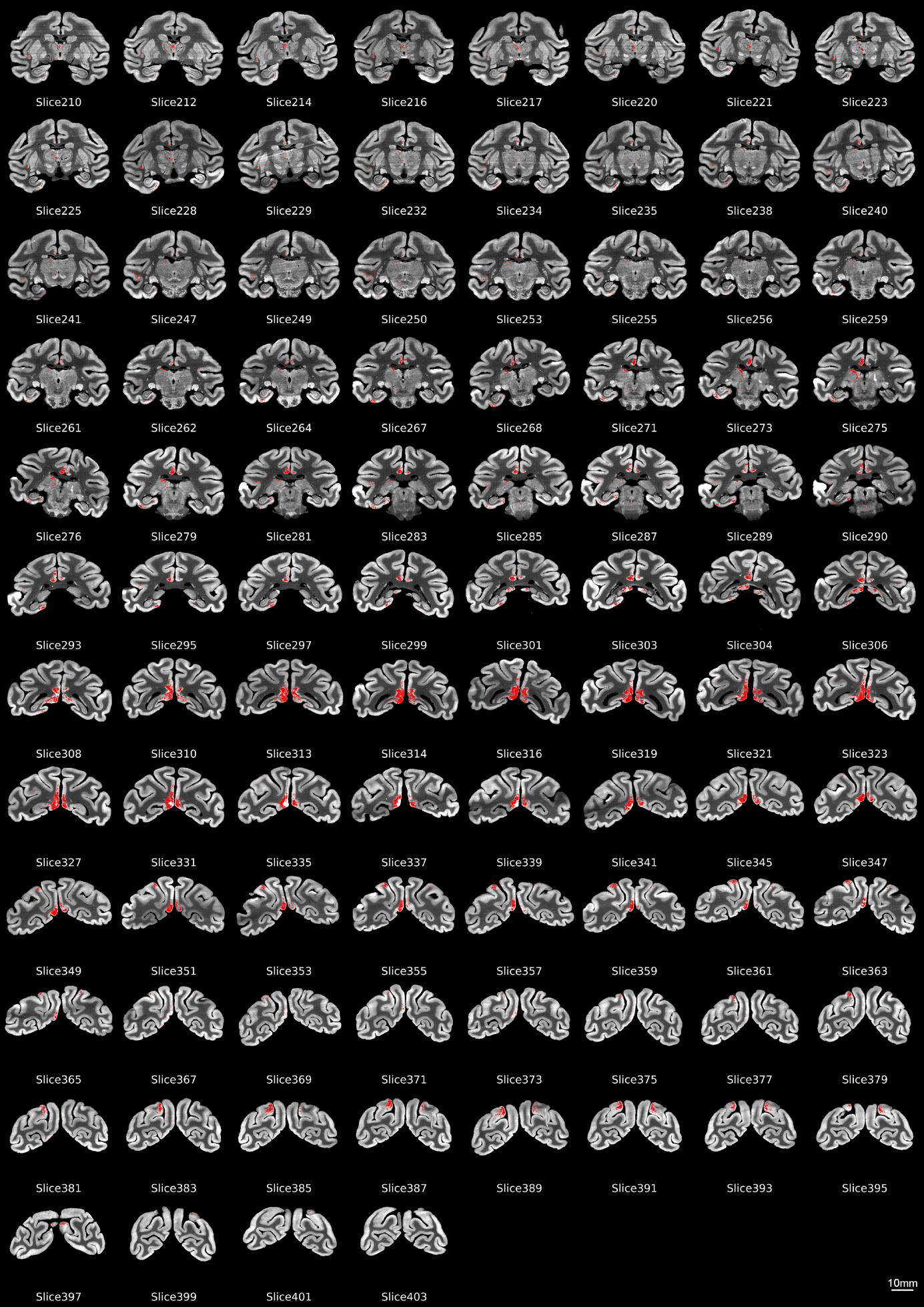

**Figure S4. Distribution of neurons projecting to area 23 in the native space. (Continued)**

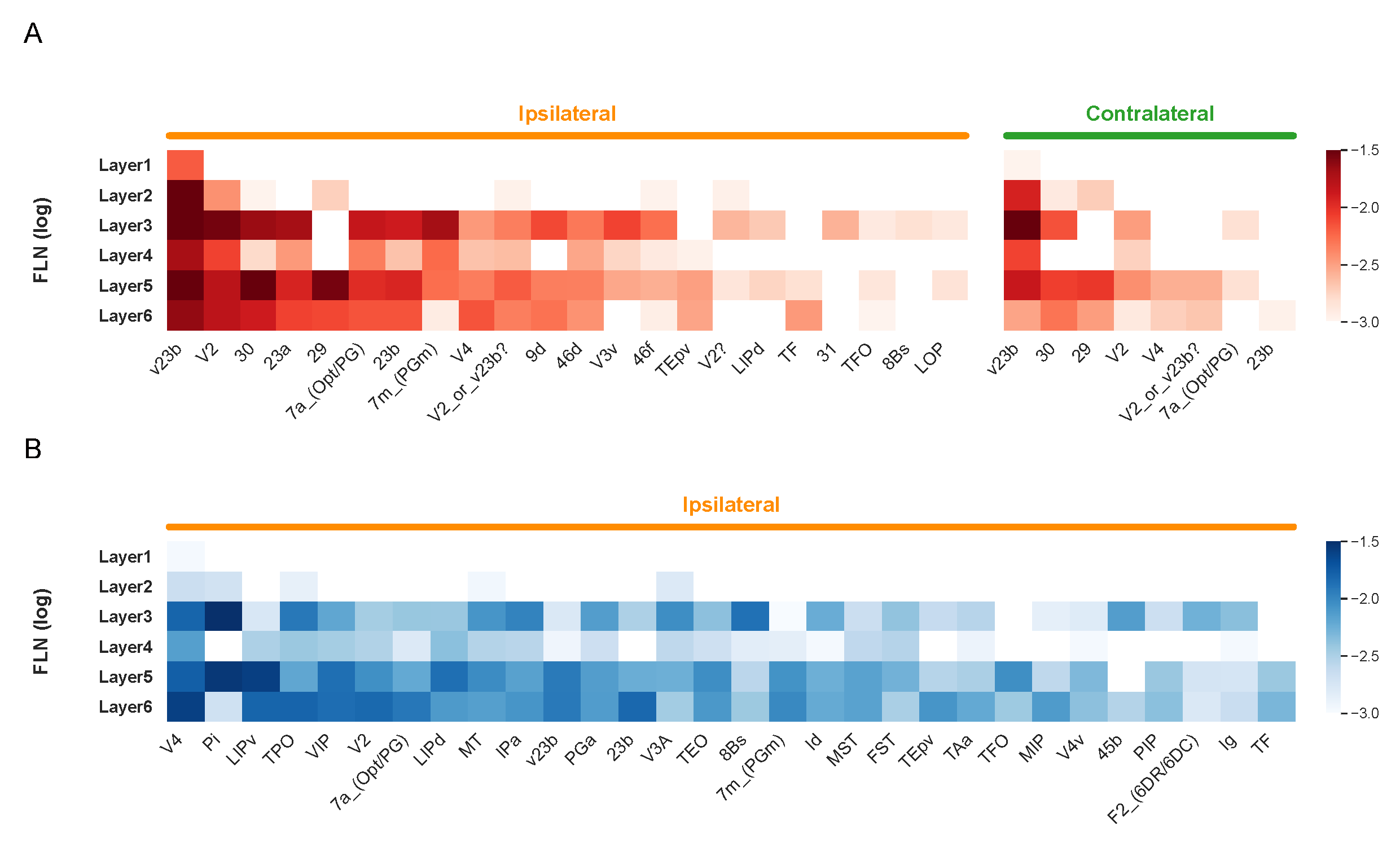

**Figure S5.** **Spatial distribution of retrograde tracing data based on the D99 143-area parcellation**

A-B Heatmaps showing the FLN of retrogradely labeled neurons (log scale) across ipsilateral and contralateral cortical areas projecting to area 23 (A) and claustrum (B).

.

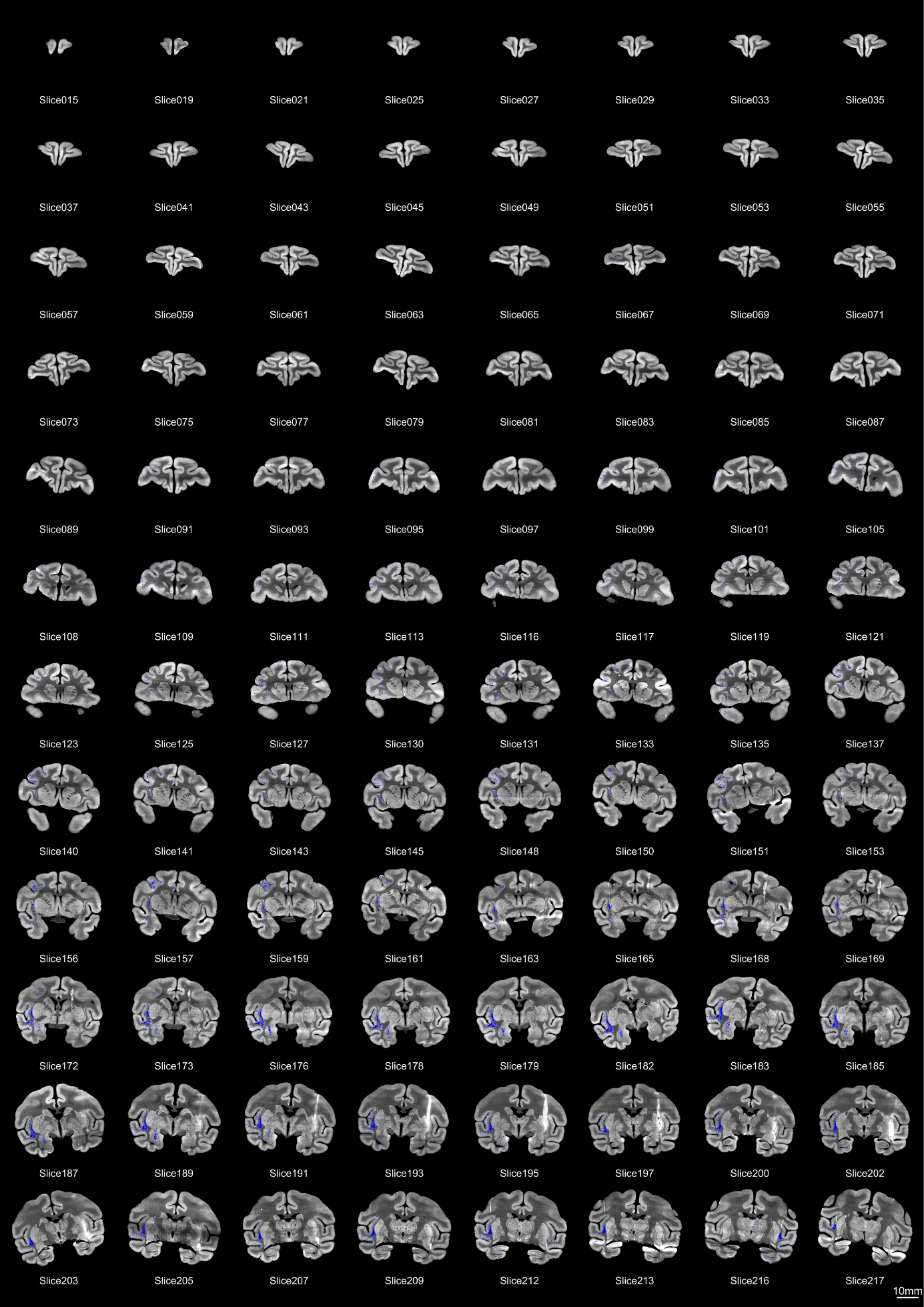

**Figure S6. Distribution of neurons projecting to claustrum in the in native space.**

Coronal sections showing the distribution of retrogradely labeled neurons in the macaque brain prior to 3D reconstruction.Blue dots represent retrogradely labeled neurons projecting to claustrum.

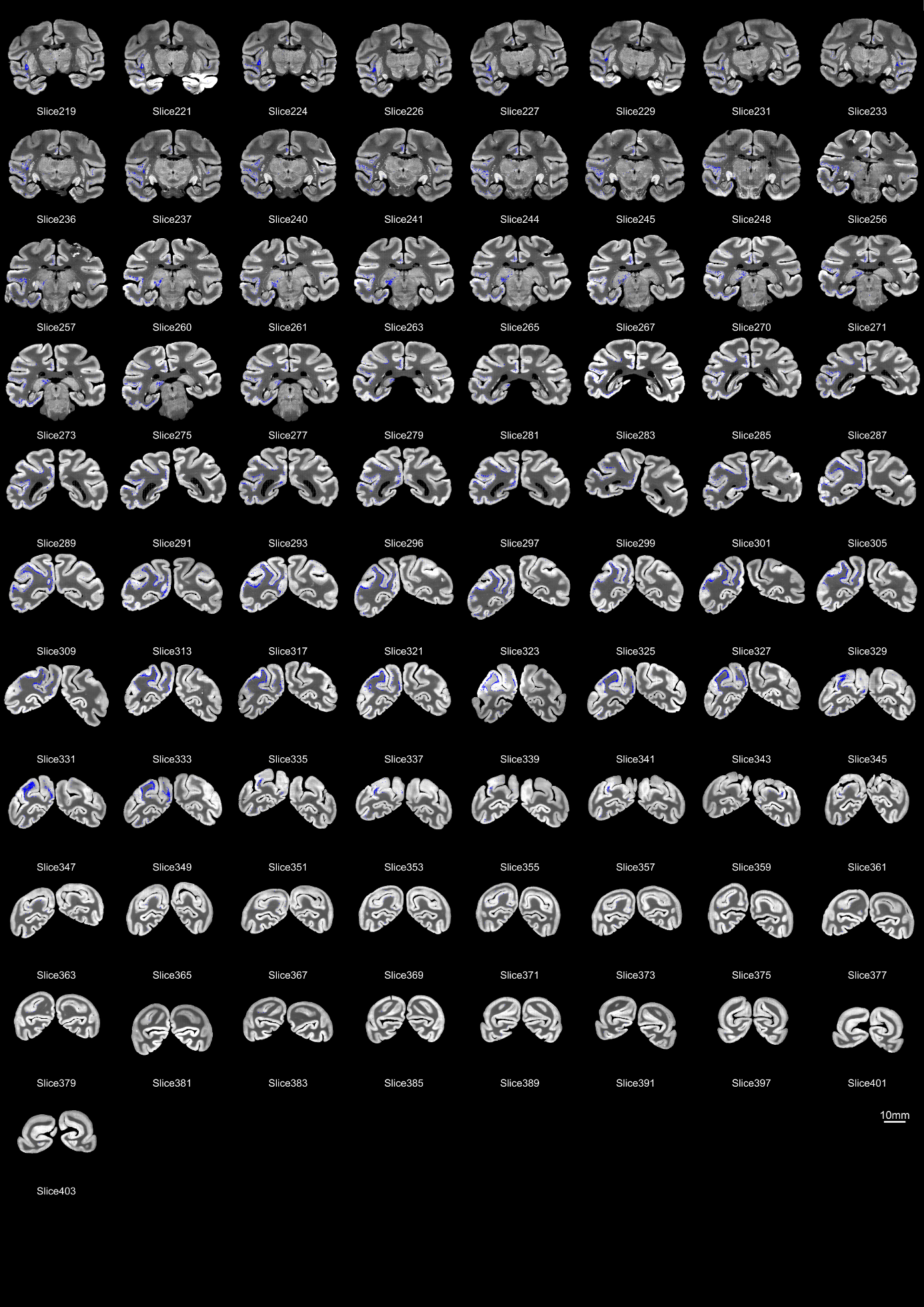

**Figure S6. Distribution of neurons projecting to claustrum in the in native space. (continued)**
